# Juvenile fluoxetine treatment affects the maturation of the medial prefrontal cortex and behavior of adolescent female rats

**DOI:** 10.1101/2024.12.11.627921

**Authors:** Joanna Kryst, Agnieszka Chocyk, Anna Solarz-Andrzejewska, Iwona Majcher-Maślanka

**Affiliations:** Department of Pharmacology and Brain Biostructure, Maj Institute of Pharmacology, Polish Academy of Sciences, Smętna Street 12, 31-343 Kraków, Poland; Laboratory of Molecular Biology, Institute of Physiotherapy and Health Sciences, Jerzy Kukuczka Academy of Physical Education, Mikołowska St. 72a, 40-065, Katowice, Poland

**Keywords:** Fluoxetine, Adolescence, Medial prefrontal cortex, Apoptosis, Endoplasmic reticulum stress, AMPA glutamate receptors

## Abstract

**Background:** Serotonin is strongly involved in the regulation of brain development. An early-life imbalance in brain serotonin levels may influence the proper formation of neuronal circuits and synaptic plasticity. One of the factors that can affect serotonin concentrations is exposure to fluoxetine (FLX), a selective serotonin reuptake inhibitor, the first-line pharmacological treatment for depression and anxiety in the pediatric population. Women are more prone to depression and anxiety from a young age. The safety of early-life FLX treatment is still questionable. We hypothesized that juvenile FLX treatment influences the brain maturation and behavior of adolescent females.

**Methods:** On postnatal days (PNDs) 20–28, juvenile female rats were injected once daily with FLX. Five days later, anxiety- and fear-related behaviors and the response to amphetamine were assessed. On PND 40, the numbers of neurons and glial cells in the medial prefrontal cortex (mPFC) and hippocampus were estimated via stereological methods. Additionally, the mRNA expression of cell survival/apoptosis and synaptic plasticity markers was evaluated via RT‒qPCR.

**Results:** Juvenile FLX attenuated anxiety-like behaviors, impaired fear memory and blunted the locomotor response to amphetamine in adolescent females. Simultaneously, FLX increased the regional volume and the numbers of neurons and astrocytes in specific subregions of the mPFC but not in the hippocampus. Additionally, FLX-treated females presented increased expression of genes regulating cell survival and reduced mRNA levels of AMPA glutamate receptors in the mPFC.

**Conclusions:** Juvenile FLX affects the maturation of the mPFC; therefore, this psychotropic drug should be used with caution in young people.

## Introduction

Serotonin (5-hydroxytryptamine, 5-HT) is a monoamine neurotransmitter and modulator involved in the regulation of a wide range of physiological and mental processes, such as cardiovascular regulation, circadian rhythms, thermoregulation, pain, eating, mood, emotion, stress response and cognitive processes (Charnay & Leger 2010, Sodhi & Sanders-Bush 2004). However, it is worth emphasizing that, prior to acting as a modulator of many essential functions in adult brain, 5-HT plays a critical role in brain development and maturation, both during embryonic and early postnatal life (Sodhi & Sanders-Bush 2004). The 5-HT network is among the earliest networks to occur during brain development and is involved in cell proliferation, differentiation, migration and programmed cell death. Moreover, 5-HT regulates dendritic growth and pruning. Thus, the 5-HT system plays a crucial role in neuronal circuit formation and synaptic plasticity (Sodhi & Sanders-Bush 2004). Clinical and animal studies have shown that manipulation of 5-HT levels may strongly affect brain development. Inactivation of the gene encoding the serotonin transporter (5-HTT) in mice (i.e., 5-HTT knockout (KO)), which elevates extracellular 5-HT levels, alters neocortical thickness, cell density (Altamura et al 2007) and apical dendritic morphology in adult animals (Wellman et al 2007). Similarly, humans carrying different variants of polymorphisms in the 5-HTT promotor region (5-HTTLPR), such as the less transcriptionally efficient allele “short” (S) vs. the more efficient allele “long” (L), present differences in gray matter density of the frontal cortex, anterior cingulate cortex and cerebellum (Canli et al 2005). Moreover, chemogenetic manipulation of 5-HT release in the prefrontal cortex (PFC) during the first weeks of postnatal life in mice affects the density and strength of excitatory synapses (Ogelman et al 2024). Accumulating data suggest that structural and functional changes induced by early-life imbalances in brain levels of 5-HT may lead to long-lasting behavioral deficits, typical for e.g., autism spectrum disorder, anxiety and mood disorders (Adjimann et al 2021, Soiza-Reilly et al 2019, Wellman et al 2007).

Many environmental conditions can influence optimal brain levels of 5-HT during development, e.g., early-life stress (Soga et al 2020), maternal diet (Ye et al 2024) or medications, such as widely prescribed antidepressants belonging to the selective serotonin reuptake inhibitors (SSRIs) group (Adjimann et al 2021, Glover & Clinton 2016). Although administering SSRIs to pregnant women is still considered safe, an increasing amount of data shows that these drugs affect brain morphometry and behavior in offspring (Glover & Clinton 2016, Koc et al 2023). For example, a recent clinical study revealed that prenatal SSRIs exposure reduced the global gray and white matter volume in children aged 7-15 years (Koc et al 2023). Moreover, numerous studies in animal models have confirmed the detrimental effects of early-life SSRIs administration on brain function. Prenatal fluoxetine (FLX), the most widely prescribed SSRI, delays brain maturation, impairs synaptic transmission and plasticity in the medial PFC (mPFC) (Bobula et al 2024) and increases anxiety- and depressive-like behaviors (Bobula et al 2024, for review, see Adjimann et al 2021, Glover & Clinton 2016). Similar effects are also observed in the case of FLX treatment during the first three postnatal weeks in rodents (for review, see Adjimann et al 2021)), which corresponds to the last weeks of gestation and infancy in humans (Ghasemi et al 2021). These observations are in line with the fact that the prenatal period and the first three weeks of life in rodents or the equivalent developmental time in humans are critical for proper brain development (Zeiss 2021). However, little is known about the impact of SSRIs exposure in midchildhood and early adolescence. During this period, massive structural and functional remodeling and refinement occur, especially in brain regions with prolonged developmental trajectories, such as the mPFC and hippocampus (HP) (Selemon 2013, Zeiss 2021). SSRIs are considered the first-line pharmacological treatment for both depression and anxiety in young people (Jacobs 2009, Rapee et al 2023, Zhou et al 2020), and these mental health problems have increased in prevalence in recent years, especially in children and adolescents (Bevilacqua et al 2023, Ludwig-Walz et al 2022). Currently, FLX is the only SSRI licensed in the majority of countries for the treatment of children from the age of 8 years (Karanges & McGregor 2011, Viswanathan 2020). However, the Food and Drug Administration (FDA) issued several “black box” warnings concerning the possible increased risk of suicidality in young people treated with SSRIs, with a recommendation to closely monitor these patients (Viswanathan 2020). Therefore, questions remain about the safety of FLX administration during the period of postnatal brain maturation. These findings prompted us to investigate the effects of repeated FLX treatment during the period of postnatal days (PNDs) 20 to 28 in rats, which corresponds to the midchildhood and early adolescence periods in humans, on selected aspects of brain maturation and the behavioral phenotype in adolescent animals (PNDs 33-40). Specifically, we studied the regional volume and number of neuronal and glial cells in the mPFC and HP. Additionally, we measured the mRNA levels of genes involved in cell survival and apoptosis (including genes implicated in the regulation of endoplasmic reticulum (ER) stress), as well as synaptic plasticity. In the behavioral part of the study, we evaluated anxiety levels, fear learning and memory and novelty- and amphetamine-induced locomotor activity in adolescents. Our study focused solely on female rats because epidemiological data clearly show that girls have a significantly greater prevalence of anxiety and depression than boys do and therefore an increased likelihood of potential exposure to SSRIs during the brain maturation period (Breslau et al 2017, Lewis et al 2020). The COVID-19 pandemic has even increased these trends (Bevilacqua et al 2023, Ludwig-Walz et al 2022).

## Materials and methods

### Animal rearing conditions

All experimental procedures were approved by the Local Ethics Committees for Animal Research in Krakow, Poland (permit no. 71/2020, issued 12/03/2020) and met the requirements of Directive 2010/63/EU of the European Parliament and the Council of September 22, 2010, on the protection of animals used for scientific purposes. All efforts were made to minimize animal suffering. Adult male and female Wistar rats were purchased from Charles River Laboratories (Sulzfeld, Germany). All animals were housed under controlled conditions with an artificial 12 h light/dark cycle (lights on from 07:00 to 19:00), 55% ± 10 humidity, and a temperature of 22 °C ± 2. Food and tap water were freely available. The rats were mated at the Maj Institute of Pharmacology, PAS, Krakow Animal Facility. The offspring of primiparous dams were used in this study. Before delivery, the dams were housed individually in standard plastic cages (38 × 24 × 19 cm). The day of birth was designated postnatal day (PND) 0. On PND 1, the litter size was standardized to ten pups per litter with the sex balance maintained, and the litters were randomly assigned to one of the following early-life treatments: juvenile FLX treatment (FLX) or juvenile vehicle treatment (VEH), i.e., the control condition.

### Juvenile FLX injections

On PNDs 20–28, juvenile female rats were injected once daily with FLX HCl (5 mg/kg/2 ml, *ip*, Sigma‒Aldrich) or with saline (2 ml/kg, *ip*) as a VEH. The FLX dose was chosen based on the data from the literature related to repeated FLX administration to juvenile rodents (Klomp et al 2012, Klomp et al 2014, Norrholm & Ouimet 2000). Moreover, an FLX dose of 5 mg/kg for rodents corresponds well with the clinically effective dose of FLX, i.e., 10–20 mg/day (approximately 0.3– 0.9 mg/kg) in adult and pediatric patients (Beasley et al 2000, Zhou et al 2020). Owing to high hepatic drug metabolism and faster elimination, FLX is administered in rodents at approximately tenfold higher doses than in humans (Caccia et al 1990, Kryst et al 2022). To minimize stress, on PNDs 20 and 21, the pups still resided with their mothers during the injection procedure. The animals were weaned on PND 22 and housed under controlled conditions in the same-litter, sex and treatment groups until PND 32. The body weights of the animals were measured daily from PND 20 to PND 28 and at the end of the experiment on PND 40 with a Kerm PCB electronic precision scale (Balingen, Germany).

### Experimental groups

A total of 31 female offspring were used in the study (16 VEH- and 15 FLX-treated rats), which originated from 3 VEH litters and 3 FLX litters. Male offspring were used in other scientific projects. On PND 32, the animals were assigned to the final experimental groups, in which a maximum of two subjects from the same litter were used. Two separate groups of animals were analyzed for i) anxiety-like behavior, fear response and mRNA expression (*n* = 7–8) and ii) novelty- and amphetamine-induced locomotor activity (*n* = 7–8) and stereological estimations of the numbers of neuronal and glial cells in the mPFC and HP (*n* = 5). The experiments were conducted after at least a 5-day FLX washout period on PNDs 33-40 (adolescence period in rats).

### Light/dark exploration test

On PND 33, 8 VEH- and 7 FLX-treated rats were subjected to the light/dark exploration test to assess anxiety-like behaviors. The light/dark exploration test was performed as described previously by Chocyk et al. (2013) and Solarz-Andrzejewska et al. (2023) (Chocyk et al 2013, Solarz-Andrzejewska et al 2023). Briefly, each experimental cage included an arena (45 × 45 × 45 cm) with a light compartment made of clear acrylic and a dark compartment made of black acrylic. The black compartment covered 33% of the total cage area, and the black dividing wall was equipped with a central tunnel gate (11 × 8.4 cm). The light compartment was brightly illuminated (100 lx), whereas the dark compartment received no light at all. The animals were kept in total darkness for 30 min prior to testing, and the entire experiment was conducted with the room lights off. The animals were individually tested in single 10 min trials. The behavioral responses during the test sessions were recorded using Fear Conditioning (FC) software (TSE, Bad Homburg, Germany). Specifically, the number of transitions between the compartments, time spent in each compartment, and locomotor activity (the distance traveled) were measured.

### Fear conditioning

On PND 37, the rats were subjected to a fear conditioning (FC) procedure. Behavioral tests were performed and analyzed via a computer-controlled FC system (TSE, Bad Homburg, Germany) as described previously by Chocyk et al. (2014) and Solarz-Andrzejewska et al. (2023) (Chocyk et al 2014, Solarz-Andrzejewska et al 2023). Each FC unit consisted of sound-attenuating housing with a loudspeaker, camera, ventilation fan and 4 symmetrically mounted lamps in the ceiling construction and test box. The test box comprised the test arena and a base construction containing integrated infrared animal detection sensors in the X, Y (horizontal) and Z (vertical) axes. The sensor frames along the axes were equipped with 32 sensor pairs mounted 14 mm apart. All sensors were scanned at a sampling rate of 100 Hz, i.e., the position of the animal was checked 100 times per second. Several pilot experiments were run that compared automatic and manual scoring of freezing behavior; the data obtained from each method were highly correlated.

During the experimental procedure, the animals were tested in two different arenas and contexts (A and B). For the first context (Context A), the arena (46 × 46 × 47 cm) was made of transparent acrylic and had a floor made up of stainless-steel rods (4 mm in diameter) spaced 8.9 mm apart (center to center). The floor was connected to a shocker-scrambler unit for delivering shocks of defined duration and intensity. The arena was cleaned with 1% acetic acid solution. A ventilation fan provided background noise (65 dB), and lamps provided uniform illumination of 60 lx inside the fear conditioning housing. During tests in Context A, the room lights remained on. The animals were subsequently transported in transparent plastic boxes. The experimenters wore white clothes and gloves.

For the second context (Context B), the arena (46 × 46 × 47 cm) was made of black acrylic (permeable to infrared light) with a gray plastic floor. The arena was cleaned with 70% ethanol solution and faintly illuminated (4 lx). The tests in Context B were conducted with the room light off. The animals were subsequently transported in black plastic boxes. The experimenters wore blue clothes and gloves.

FC and memory were assessed via the Pavlovian paradigm. On Day 1 of the experiment, the animals were subjected to the FC procedure in Context A (acquisition/training). The animals were placed in Context A and allowed to habituate for 180 s. Next, the animals received five tone‒ shock pairings in which the tone (amplitude: 80 dB; frequency: 2 kHz; duration: 10 s) was coterminated with foot shock (intensity: 0.7 mA; duration: 1 s). The intertrial interval was 60 s. Animals were removed from Context A 60 s after the last trial.

On Day 2, all the animals were once again exposed to Context A and were left undisturbed for 6 min (expression of contextual fear conditioning (CFC)) and then returned to their home cages. Two hours later, the animals were placed in a new context (Context B) and, after 180 s of habituation, received five tone-alone presentations with 61 s intertrial intervals (expression of auditory fear conditioning (AFC)). The animals were removed from Context B 60 s after the last trial. Behavioral responses during all the sessions were recorded and automatically analyzed via FC software (TSE, Bad Homburg, Germany). Freezing (i.e., immobility) was taken as the behavioral measure of fear and was defined as the absence of all nonrespiratory movements for at least 2 s. The cumulative duration of freezing was calculated for each session and expressed as a percentage of the entire session time, excluding habituation time.

### Novelty- and amphetamine-induced locomotor activity tests

On PND 33, a separate group of animals (8 VEH- and 7 FLX-treated females) was subjected to the locomotor activity test. Locomotor activity was recorded and analyzed individually for each animal via Opto-Varimex cages (43 × 44 cm) and AutoTrack software (Columbus Instruments, OH, USA), as described previously by Majcher-Maślanka et al. (2019) and Solarz-Andrzejewska et al. (2023) (Majcher-Maslanka et al 2019, Solarz-Andrzejewska et al 2023). The rats were placed in test cages without previous habituation and were allowed to explore the environment freely for 20 min (novelty-induced locomotion, session 1). Next, the rats received vehicle injections (saline, 1 ml/kg *sc*) and were left in test cages for 50 min (session 2). Finally, the same rats were injected with amphetamine (1 mg/kg) and left in test cages for another 50 min (session 3). The locomotor activity of the animals was recorded for each session separately. The data are presented as the average distance traveled over the entire session.

### Cresyl violet staining and immunohistochemistry

Nissl (cresyl violet, CV) staining and immunohistochemistry were performed as previously described by Chocyk et al. (2011) and Majcher-Maślanka et al. (2019) (Chocyk et al 2011, Majcher-Maslanka et al 2019). Briefly, on PND 40, 5 VEH- and 5 FLX-treated females were deeply anesthetized and transcardially perfused with saline followed by 4% paraformaldehyde in 0.1 M PBS (pH 7.4). The brains were removed from the skulls, postfixed in 4% paraformaldehyde in PBS for 24 h at 4 °C, and sectioned using a vibratome (VT1000S, Leica, Wetzlar, Germany) into 50-μm thick coronal slices at the level of the mPFC (Bregma +3.70 to +2.20 mm) and the HP (Bregma −1.8 to −6.3) according to a stereotaxic atlas of the rat brain (Paxinos and Watson, 1998). Every fourth (for the mPFC) or seventh section (for the HP) was preserved for further processing (∼10 sections from each subject).

For CV staining, the sections were mounted onto gelatin-coated slides and air-dried. Then, the slides were soaked in a series of descending alcohol solutions and submerged in a 0.5% CV acetate solution. Finally, all the sections were dehydrated and coverslipped with Permount mounting medium (Fisher Scientific).

For immunohistochemical staining of specific glial cell markers, free-floating sections were washed in 0.01 M PBS (pH 7.4) and incubated for 30 min in PBS containing 0.3% H_2_O_2_ and 0.2% Triton X-100. The sections were then rinsed and transferred to a blocking buffer (5% solution of appropriate normal serum (goat or rabbit) in 0.2% Triton X-100 in PBS) for 1 h. After the blocking procedure, the sections were incubated for 48 h at 4 °C with a goat anti-glial fibrillary acidic protein (GFAP) antibody (1:300; Santa Cruz) or a rabbit anti-ionized calcium-binding adapter molecule/allograft inflammatory factor 1 (IBA1/AIF-1) antibody (1:1,500; Proteintech). The antibodies were diluted with 3% normal serum and 0.2% Triton X-100 in PBS. After being washed in PBS, the sections were incubated for 1 h with an appropriate solution of biotinylated secondary antibodies (goat anti-rabbit or rabbit anti-goat IgG, 1:500; Vector Laboratories), followed by a 1 h incubation with an avidin-biotin-peroxidase complex (1:200, 1 h; Vectastain ABC Kit, Vector Laboratories). The immunochemical reaction was developed in a diaminobenzidine (DAB)-nickel solution containing 0.02% DAB, 0.01% H_2_O_2_ and 0.06% NiCl_2_ in TBS, which stained the immunoreactive material black. The sections were mounted onto gelatin-coated slides, air-dried and coverslipped using Permount (Fisher Scientific) as the mounting medium.

### Cell counting

The number of CV-stained or immunoreactive (IR) cells was estimated via unbiased stereological methods (West et al 1991), as described previously by Chocyk et al. (2011) and Majcher-Maślanka et al. (2019) (Chocyk et al 2011, Majcher-Maslanka et al 2019). Briefly, every fourth (for the mPFC) or seventh section (for the HP), selected by systematic random sampling along the rostrocaudal axis of the brain, was chosen for analysis (10 sections per animal). Optical fractionator sampling was performed using a Leica DM 6000 B microscope equipped with a motorized stage (Ludl Electronic Products, Hawthorne, NY, USA) connected to a controller (MAC 5000, Ludl) and a digital camera (MBF C×9000, Williston, VT, USA). Sampling was performed using Stereo Investigator 8.0 software (MBF Bioscience, Williston, VT, USA) by experimenters unaware of treatment group allocation to the specific slides. The studied regions of the mPFC, i.e., the cingulate cortex 1 (Cg1), prelimbic cortex (PL) and infralimbic cortex (IL), or the HP, i.e., CA1, CA2-CA3 and dentate gyrus (DG), were outlined under low magnification (2.5x) according to a stereotaxic atlas of the rat brain (Paxinos & Watson 1998). Sampling was performed bilaterally under high magnification (63x, oil-immersion objective) using counting frames with areas of 2,500 µm^2^ and heights of 15 µm.

Strict histological and morphological criteria were employed to discriminate between neuronal and nonneuronal CV-stained cells during the counting procedure, as described previously in detail by Chocyk et al. (2011) and Majcher-Maślanka et al. (2019). The total number of CV-positive neurons or IR glial cells per region was estimated from the number of cells sampled within the optical dissectors and calculated by multiplying the numerical density of the cells (the number of cells/mm^3^) by the regional volume occupied by the cells within the studied region. The regional volumes of the studied brain areas in each animal were determined via the Cavalieri method (West 2012). The final results are presented as the estimated total number of cells within the specific regions.

### RT‒qPCR

The RT‒qPCR procedure was performed as described previously by Solarz et al. (2021, 2023) (Solarz-Andrzejewska et al 2023, Solarz et al 2021a). Briefly, on PND 40, 6 VEH- and 6 FLX-treated females were sacrificed by decapitation, and the brains were immediately removed from the skulls. The mPFC (including the Cg1, PL, and IL regions) was dissected from 1 mm thick coronal sections using a rodent brain matrix (Ted Pella Inc., CA, USA), whereas the whole HP was dissected freehand. After dissection, the brain tissue was quickly frozen in liquid nitrogen and stored at −80 °C for later use. Total RNA from the brain tissue was extracted using the RNeasy Mini Kit (Qiagen). The total RNA concentration was measured using an Eon absorbance microplate reader and Gen 5 software (BioTek, Winooski, VT, USA). The RNA was reverse transcribed using a High-Capacity cDNA Reverse Transcription Kit (Thermo Fisher Scientific, MA). Quantitative real-time PCR was performed in duplicate with TaqMan® Gene Expression Assays (Thermo Fisher Scientific, MA; Table 1) using TaqMan™ Universal Master Mix II, no UNG (Thermo Fisher Scientific, MA) and the QuantStudio 12K Flex System (Thermo Fisher Scientific, MA). Real-time PCR was conducted under the following conditions: 50 °C for 2 min and 95 °C for 10 min, followed by 40 cycles of 95 °C for 15 s and 60 °C for 1 min. The abundance of RNA was calculated according to the following equation: abundance = 2^−(threshold^ ^cycle)^ (Livak & Schmittgen 2001). The results were normalized to glyceraldehyde-3-phosphate dehydrogenase (Gapdh) expression levels.

**Table 1.**
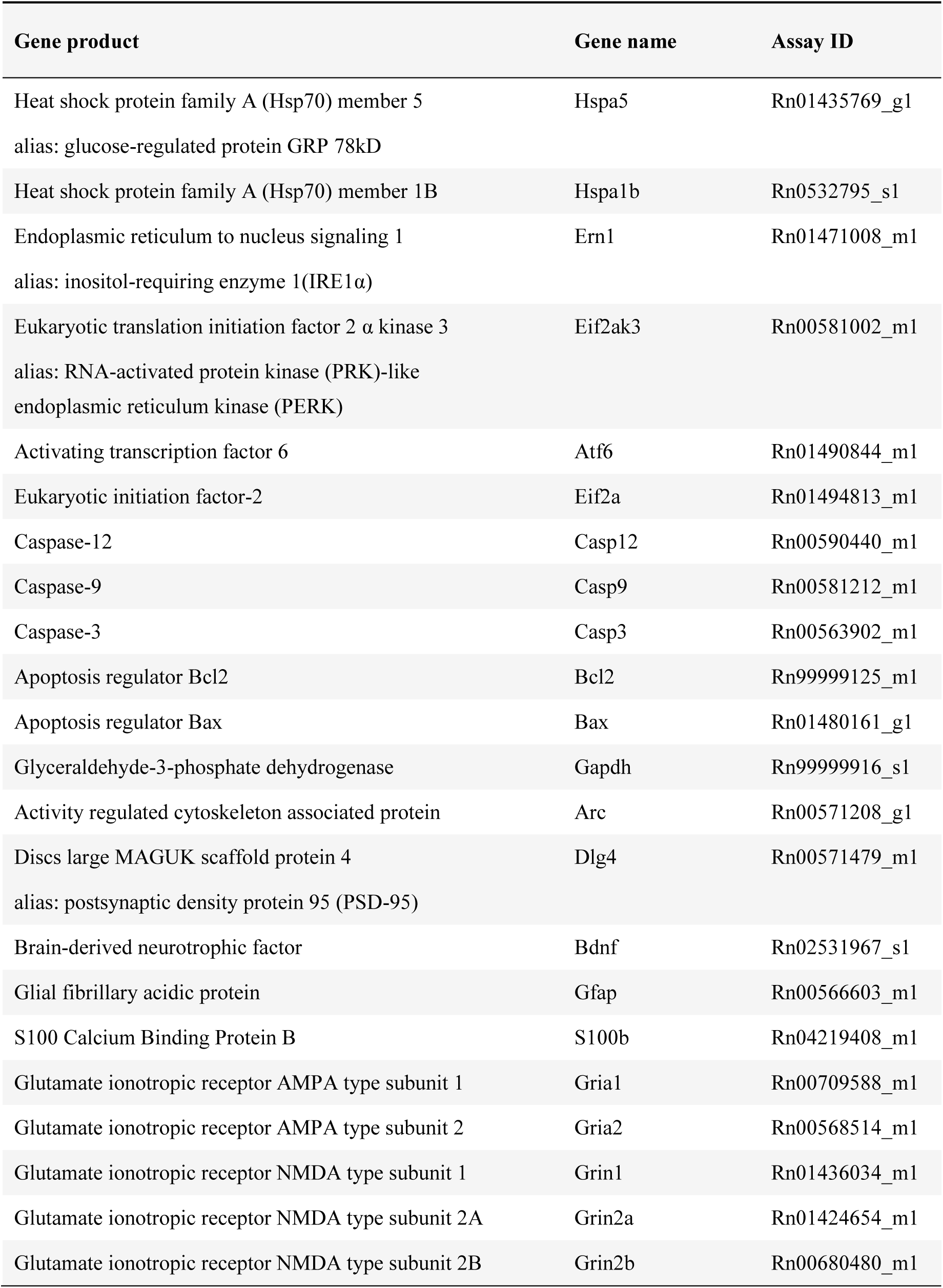
The list of TaqMan® Gene Expression Assays used in the study.

| Gene product | Gene name | Assay ID |
| --- | --- | --- |
| Heat shock protein family A (Hsp70) member 5<br>alias: glucose-regulated protein GRP 78kD | Hspa5 | Rn01435769_g1 |
| Heat shock protein family A (Hsp70) member 1B | Hspa1b | Rn0532795_s1 |
| Endoplasmic reticulum to nucleus signaling 1<br>alias: inositol-requiring enzyme 1(IRE1α) | Ern1 | Rn01471008_m1 |
| Eukaryotic translation initiation factor 2 $\alpha$ kinase 3<br>alias: RNA-activated protein kinase (PRK)-like<br>endoplasmic reticulum kinase (PERK) | Eif2ak3 | Rn00581002_m1 |
| Activating transcription factor 6 | Atf6 | Rn01490844_m1 |
| Eukaryotic initiation factor-2 | Eif2a | Rn01494813_m1 |
| Caspase-12 | Casp12 | Rn00590440_m1 |
| Caspase-9 | Casp9 | Rn00581212_m1 |
| Caspase-3 | Casp3 | Rn00563902_m1 |
| Apoptosis regulator Bcl2 | Bcl2 | Rn99999125_m1 |
| Apoptosis regulator Bax | Bax | Rn01480161_g1 |
| Glyceraldehyde-3-phosphate dehydrogenase | Gapdh | Rn99999916_s1 |
| Activity regulated cytoskeleton associated protein | Arc | Rn00571208_g1 |
| Discs large MAGUK scaffold protein 4<br>alias: postsynaptic density protein 95 (PSD-95) | Dlg4 | Rn00571479_m1 |
| Brain-derived neurotrophic factor | Bdnf | Rn02531967_s1 |
| Glial fibrillary acidic protein | Gfap | Rn00566603_m1 |
| S100 Calcium Binding Protein B | S100b | Rn04219408_m1 |
| Glutamate ionotropic receptor AMPA type subunit 1 | Gria1 | Rn00709588_m1 |
| Glutamate ionotropic receptor AMPA type subunit 2 | Gria2 | Rn00568514_m1 |
| Glutamate ionotropic receptor NMDA type subunit 1 | Grin1 | Rn01436034_m1 |
| Glutamate ionotropic receptor NMDA type subunit 2A | Grin2a | Rn01424654_m1 |
| Glutamate ionotropic receptor NMDA type subunit 2B | Grin2b | Rn00680480_m1 |

### Statistical analysis

Statistical analysis of the data was performed using Statistica 13.3 software (TIBCO Software Inc., USA). Initially, the data were tested for a normal distribution and homogeneity of variances using Shapiro-Wilk test and Levene’s test, respectively. Data that followed a normal distribution and had equal variances among groups were further analyzed by Student’s t test (for comparisons between the VEH and FLX groups). Amphetamine-induced locomotor activity was specifically analyzed via mixed-design ANOVA, with early-life VEH/FLX treatment as a between-subjects factor and saline and amphetamine injections as within-subjects factors. Data that did not show a normal distribution or homogeneity of variance were analyzed via the Mann–Whitney U test (for comparisons between the VEH and FLX groups). *P* values < 0.05 were considered significantly different.

## Results

### Effects of juvenile FLX treatment on body weight in female rats

Owing to the potential anorectic effects of FLX, the body weights of juvenile female rats were carefully monitored during the injection period (PND 20-28) and also measured at the end of the experiment on PND 40. The differences in body weight between PNDs 28–20 and PNDs 40–20 were subsequently calculated (body weight gain indices) (Fig. 1A, B). Statistical analysis revealed no significant effects of FLX on body weight gain in juvenile and adolescent females (PND 28-20: *U*_16,15_ = 74.5, *p* = 0.075, Mann–Whitney U test; PND 40-20: *t*_29_ = 0.950, *p* = 0.350, Student’s t test) (Fig. 1A, B).

**Fig. 1.**
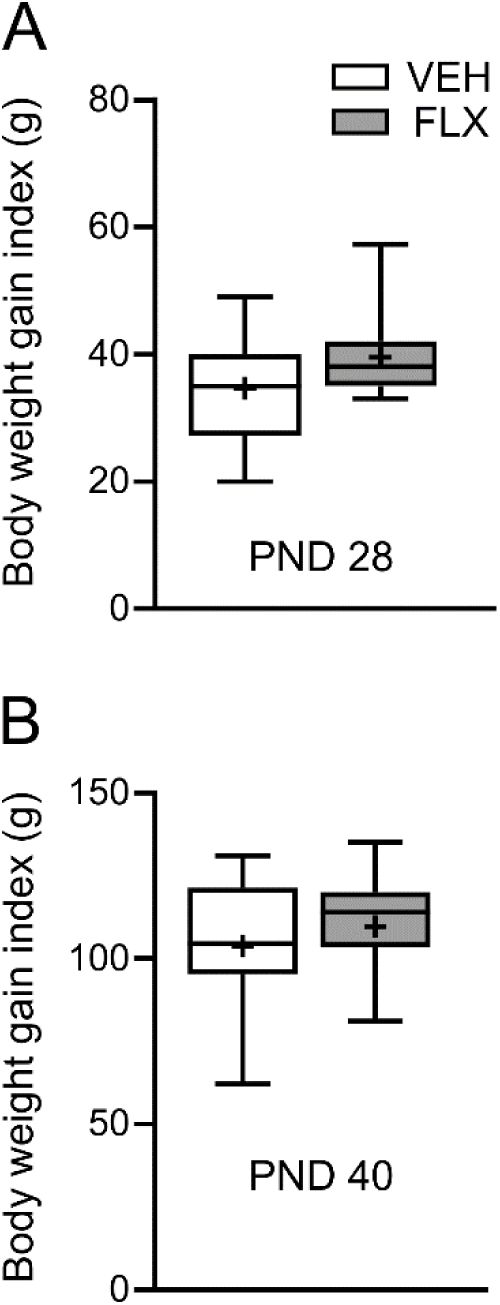
Effects of juvenile FLX treatment on body weight gain in female rats. The data are presented as box plots and expressed as the body weight gain index calculated as the body weight difference between PND 28 and PND 20 (A) and between PND 40 and PND 20 (B). In the box plots, the boxes depict the 25th and 75th quartiles, the horizontal line and plus sign correspond to the median and the mean, respectively, whereas the whiskers point to the minimum and maximum of the data (*n* = 15–16). The Mann–Whitney U test (A) and Student’s t test (B) did not reveal any significant differences between the experimental groups. Abbreviations: FLX, fluoxetine; PND, postnatal day; VEH, vehicle.

### Behavioral effects of juvenile FLX treatment in adolescent females

After a 5-day FLX washout period, on PNDs 33-40, adolescent females were subjected to behavioral tests to assess anxiety-like behaviors, fear memory and novelty- and amphetamine-induced locomotor activity.

Anxiety-like behaviors were evaluated in the light/dark box test. Statistical analysis revealed that FLX-treated females had lower levels of anxiety than control females did (Fig. 2). Specifically, juvenile FLX treatment increased the distance traveled by adolescent females on the light side (*t*_13_ = 2.31, *p* = 0.038) (Fig. 2A) and the time spent in the light compartment (*t*_13_ = 2.71, *p* = 0.018, Student’s t test) (Fig. 2B). Compared with VEH-treated females, FLX-treated females also presented a greater number of transitions between the light and dark compartments (*t*_13_ = 3.29, *p* = 0.006; Student’s t test) (Fig. 2C). Concurrently, FLX did not affect the total distance traveled during the test (VEH: 7,896.2 ± 1,954.1, FLX: 8,863.5 ± 1,779.0; *t*_13_ = 0.997, *p* = 0.334; Student’s t test).

**Fig. 2.**
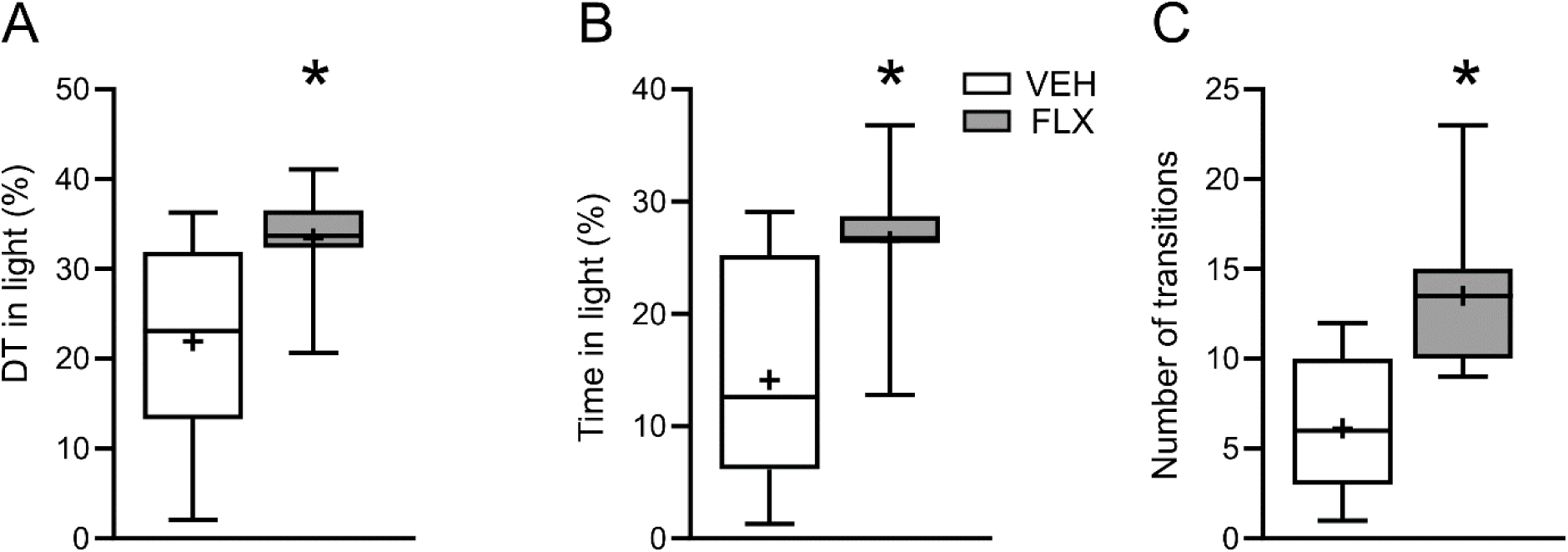
Effects of juvenile FLX treatment on the anxiety-like behavior of adolescent females in the light/dark exploration test. The intensity of anxiety-like behavior was assessed as the distance traveled on the light side (A), the time spent on the light side (B), and the number of transitions between the light and dark sides (C). The data are presented as box plots. In the box plots, the boxes depict the 25th and 75th quartiles, the horizontal line and plus sign correspond to the median and the mean, respectively, whereas the whiskers point to the minimum and maximum of the data (*n* = 7–8). \**p* < 0.05 compared with VEH rats (Student’s t test). Abbreviations: DT, distance traveled; FLX, fluoxetine; VEH, vehicle.

The acquisition and expression of fear memory were studied in the FC paradigm. Compared with control rats, FLX-treated females presented impaired FC, manifested as reduced freezing behavior during the FC training session (Day 1, context A) (*t*_13_ = 3.03, *p* = 0.01, Student’s t test) (Fig. 3A). Moreover, juvenile FLX treatment also affected the expression of both the CFC (*U*_8,7_ = 9.0, *p* = 0.032, Mann–Whitney U test) (Fig. 3B) and the AFC (*t*_13_ = 2.70, *p* = 0.018, Student’s t test) (Fig. 3C). Specifically, on Day 2 of the experiment, FLX-treated females presented a lower level of freezing behavior in response to both contexts A and B than VEH-treated females did (Fig. 3B, C). The analysis of locomotor activity during the specific sessions of the FC test revealed that FLX did not affect the total distance traveled during the acquisition session on Day 1 of the experiment (VEH: 5,508.21 ± 1,338.71, FLX: 6,543.03 ± 842.57; *t*_13_ = 1.759, *p* = 0.102, Student’s t test). However, FLX-treated females showed greater distance traveled during both the CFC and AFC expression sessions on Day 2 (CFC: VEH: 1,363.89 ± 805.15, FLX: 2,962.07 ± 1,835.32; *t*_13_ = 2.238, *p* = 0.043, Student’s t test; AFC: VEH: 4,551.10 ± 617.50, FLX: 6,765.51 ± 2,248.25; *U*_8,7_ = 9.0, *p* = 0.032, Mann–Whitney U test).

**Fig. 3.**
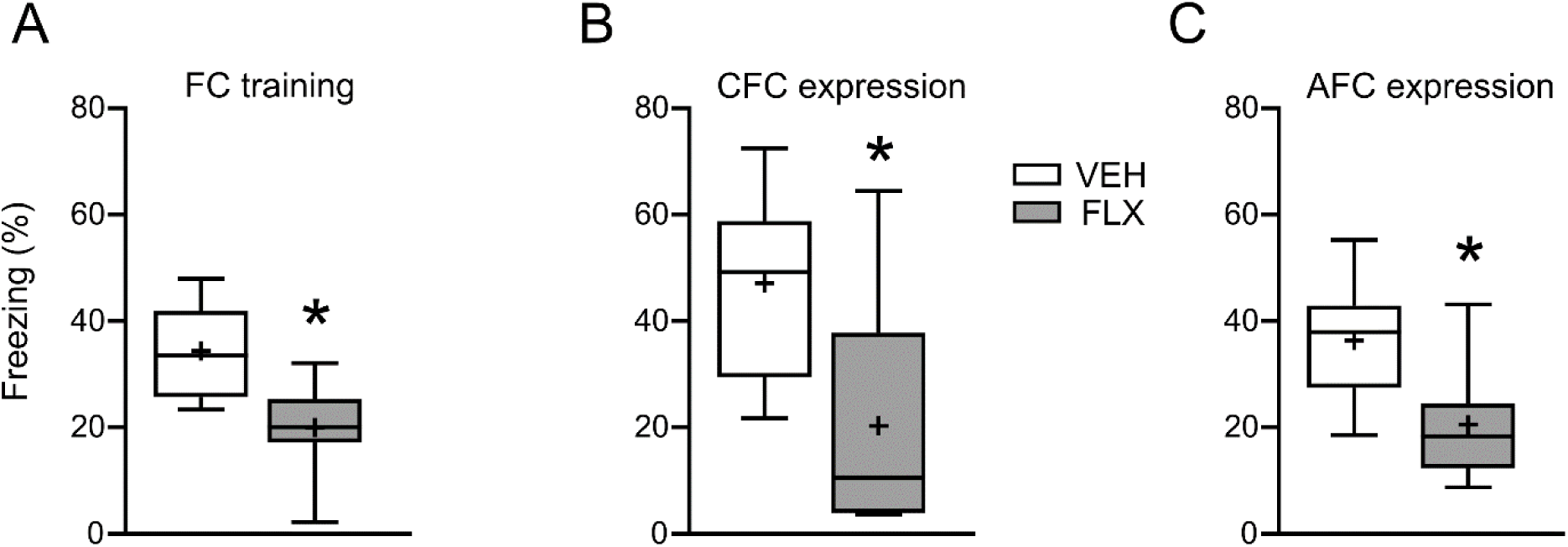
Effects of juvenile FLX treatment on the acquisition (A) and expression (B, C) of conditioned fear in adolescent females. The data are presented as box plots and expressed as freezing behavior in a percentage of the session time. In the box plots, the boxes depict the 25th and 75th quartiles, the horizontal line and plus sign correspond to the median and the mean, respectively, whereas the whiskers point to the minimum and maximum of the data (*n* = 7–8). \**p* < 0.05 compared with VEH rats (Student’s t test in A, C and the Mann–Whitney U test in B). Abbreviations: AFC, auditory fear conditioning; CFC, contextual fear conditioning; FC, fear conditioning; FLX, fluoxetine; VEH, vehicle.

The analysis aimed to specifically measured the locomotor activity of the animals in standard activity cages (Opto-Varimex) revealed that, in response to the novel environment of the cage, adolescent female behavior did not differ between the treatment groups (*t*_13_ = 0.348, *p* = 0.733, Student’s t test) (Fig. 4A). However, when acute saline and amphetamine injections were administered in a familiar environment, juvenile FLX-treated females generally presented lower locomotor activity than control females did, regardless of the type of injection (juvenile VEH/FLX treatment as between-subjects factor: *F*_1,13_ = 4.89, *p* = 0.046; juvenile treatment × saline/amphetamine injection: *F*_1,13_ = 2.10, *p* = 0.171; mixed-design ANOVA) (Fig. 4B). Nonetheless, both main experimental groups responded to acute amphetamine treatment with enhanced locomotor activity (saline/amphetamine injection as within-subjects factor: *F*_1,13_ = 42.08, *p* < 0.0001, mixed-design ANOVA) (Fig. 4B).

**Fig. 4.**
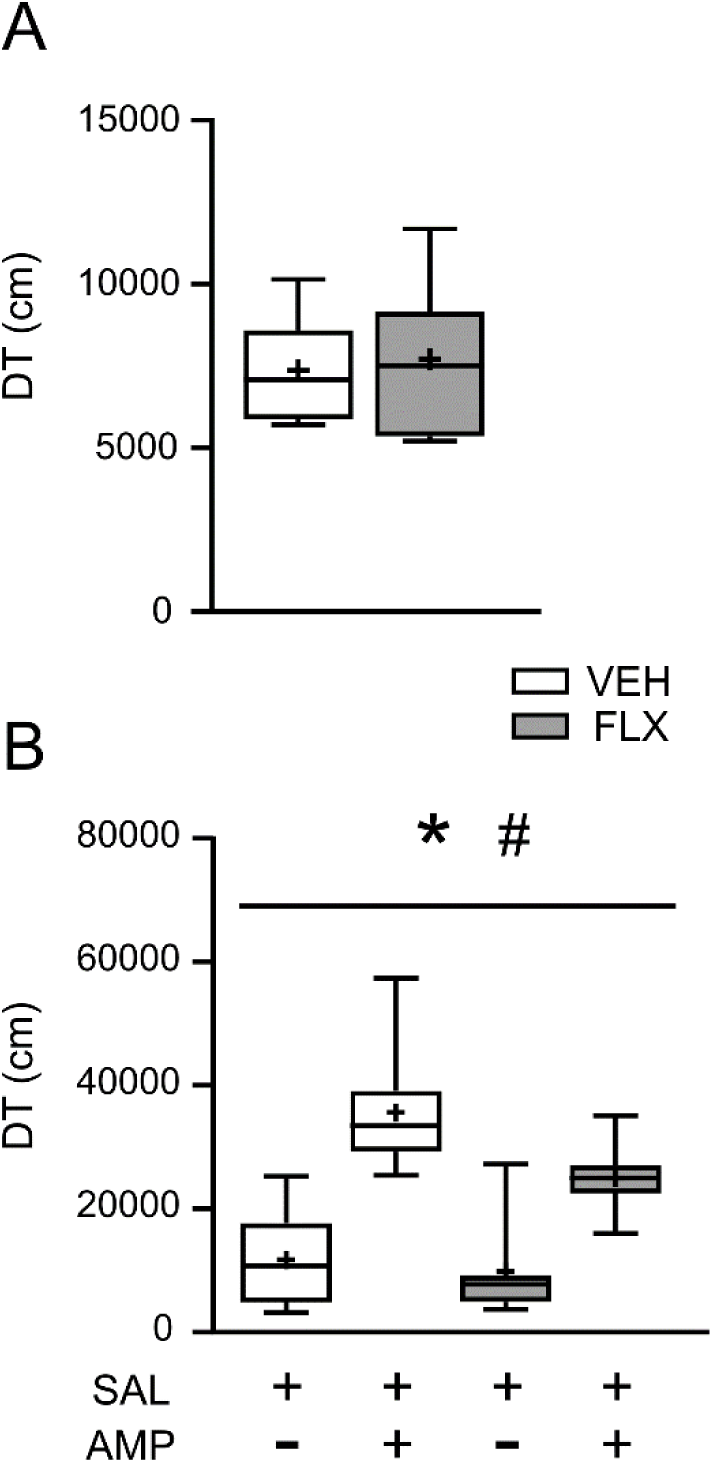
Effects of juvenile FLX treatment on locomotor activity in adolescent females. (A) Novelty-induced locomotion, (B) amphetamine-induced locomotor activity. The data are presented as box plots and expressed as the total distance traveled during a specific session. In the box plots, the boxes depict the 25th and 75th quartiles, the horizontal line and plus sign correspond to the median and the mean, respectively, whereas the whiskers point to the minimum and maximum of the data (*n* = 7–8). \**p* < 0.05 – the effect of juvenile FLX treatment, #*p* < 0.05 – the effect of amphetamine injection in adolescence as within-subjects factor (mixed-design ANOVA). Abbreviations: AMP, amphetamine; DT, distance traveled; FLX, fluoxetine; SAL, saline; VEH, vehicle.

### Effects of juvenile FLX treatment on regional volume and the numbers of neurons and glial cells in the mPFC and HP of adolescent females

Stereological estimations of the volume and number of specific populations of cells in the mPFC revealed that juvenile FLX treatment affected many of the studied parameters. First, FLX-treated females presented greater mPFC volume in each of its regions (Cg1: *t*_8_ = 3.26, *p* = 0.011, Student’s t test; PL: *U*_5,5_ = 0.0, *p* = 0.012; IL: *U*_5,5_ = 0.0, *p* = 0.012, Mann–Whitney U test) (Fig. 5A). Concurrently, FLX did not influence the numerical density of specific cells in the mPFC (ESM_1: Table S1). However, when the total number of cells per regional volume was estimated, females injected with FLX during juvenility presented more neuronal cells in the Cg1 (*t*_8_ = 2.36, *p* = 0.046) and PL (*t*_8_ = 3.09, *p* = 0.015, Student’s t test) than VEH-treated females did (Fig. 5B). Additionally, juvenile FLX treatment increased the total number of GFAP-IR glial cells specifically in the IL region of the mPFC (IL: *t*_8_ = 3.29, *p* = 0.011; Cg1: *t*_8_ = 0.104, *p* = 0.920; PL: *t*_8_ = 0.656, *p* = 0.530; Student’s t test) (Fig. 6A). There was no statistically significant effect of FLX on the total number of IBA1-IR microglia in the mPFC of adolescent females (Cg1: *t*_8_ = 1.03, *p* = 0.333; PL: *t*_8_ = 0.156, *p* = 0.880; IL: *t*_8_ = 2.21, *p* = 0.058; Student’s t test) (Fig. 6B). However, a trend toward a lower number of microglia in the IL of FLX-treated females was clearly observed (*p* = 0.058). Representative photomicrographs showing examples of CV histological staining and GFAP and IBA1 immunostaining in the mPFC of adolescent females are presented in Fig. 7.

**Fig. 5.**
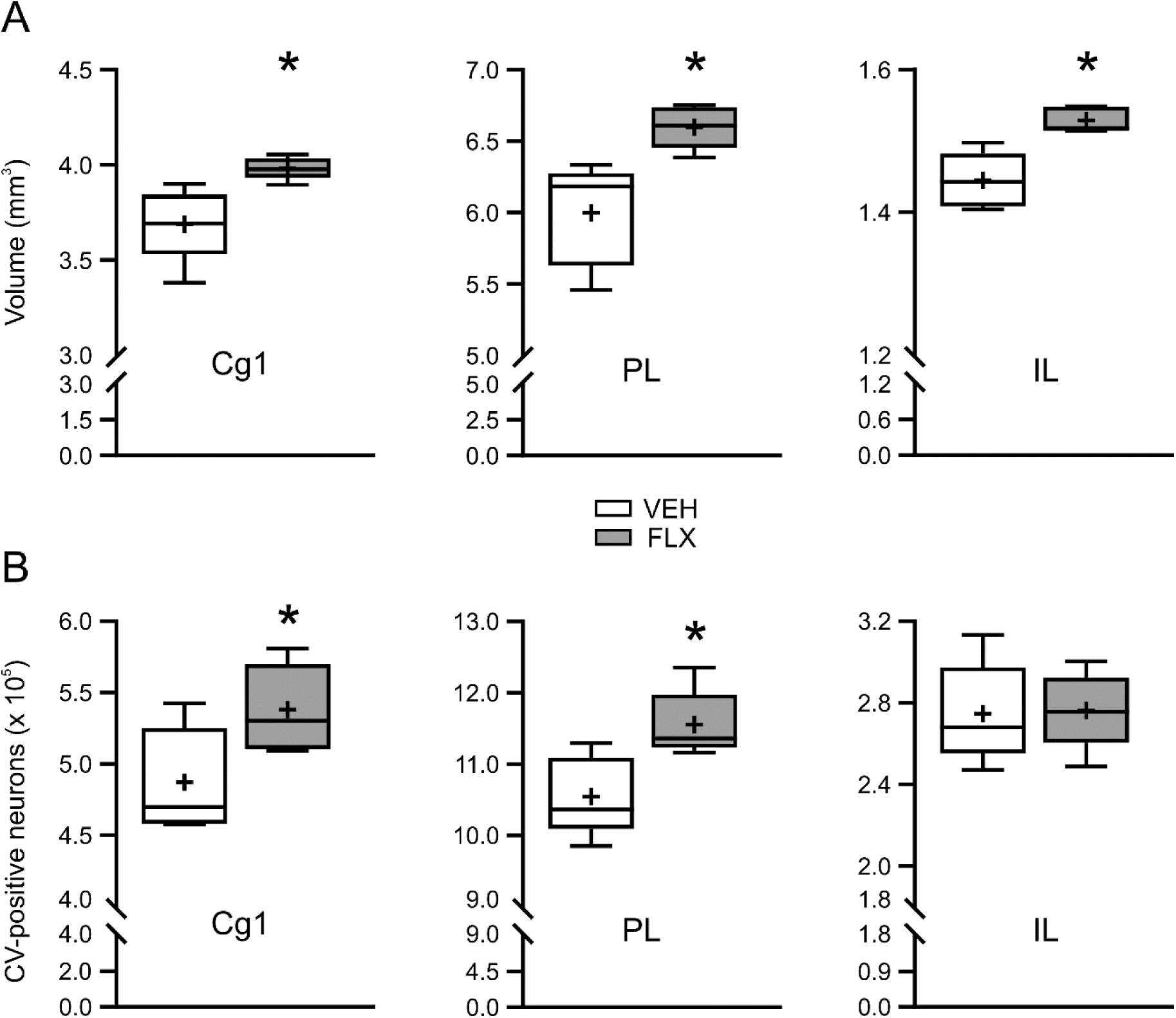
Effects of juvenile FLX treatment on regional volume and the number of neurons in the mPFC of adolescent females. The data are presented as box plots. In the box plots, the boxes depict the 25th and 75th quartiles, the horizontal line and plus sign correspond to the median and the mean, respectively, whereas the whiskers point to the minimum and maximum of the data (*n* = 5). \**p* < 0.05 compared with VEH rats (Student’s t test in A (Cg1), B and the Mann–Whitney U test in A (PL, IL)). Abbreviations: Cg1, cingulate cortex 1; CV, cresyl violet; FLX, fluoxetine; IL, infralimbic cortex; PL, prelimbic cortex; VEH, vehicle.

**Fig. 6.**
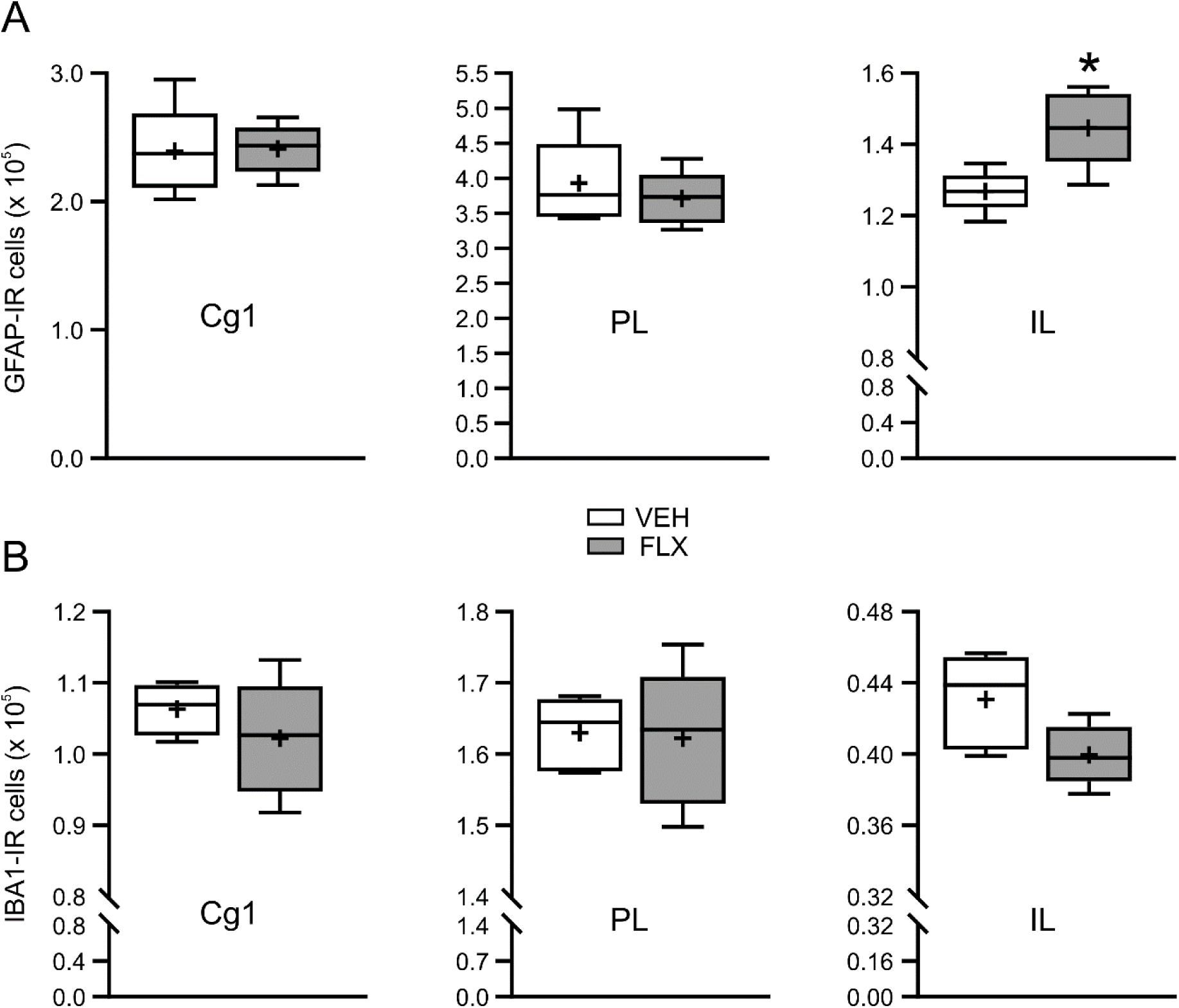
Effects of juvenile FLX treatment on the number of GFAP-IR astrocytes (A) and IBA1-IR microglia (B) in the mPFC of adolescent females. The data are presented as box plots. In the box plots, the boxes depict the 25th and 75th quartiles, the horizontal line and plus sign correspond to the median and the mean, respectively, whereas the whiskers point to the minimum and maximum of the data (*n* = 5). \**p* < 0.05 compared with VEH rats (Student’s t test). Abbreviations: Cg1, cingulate cortex 1; FLX, fluoxetine; GFAP, glial fibrillary acidic protein; IBA1, ionized calcium-binding adaptor molecule 1; IL, infralimbic cortex; IR, immunoreactive; PL, prelimbic cortex; VEH, vehicle.

**Fig. 7.**
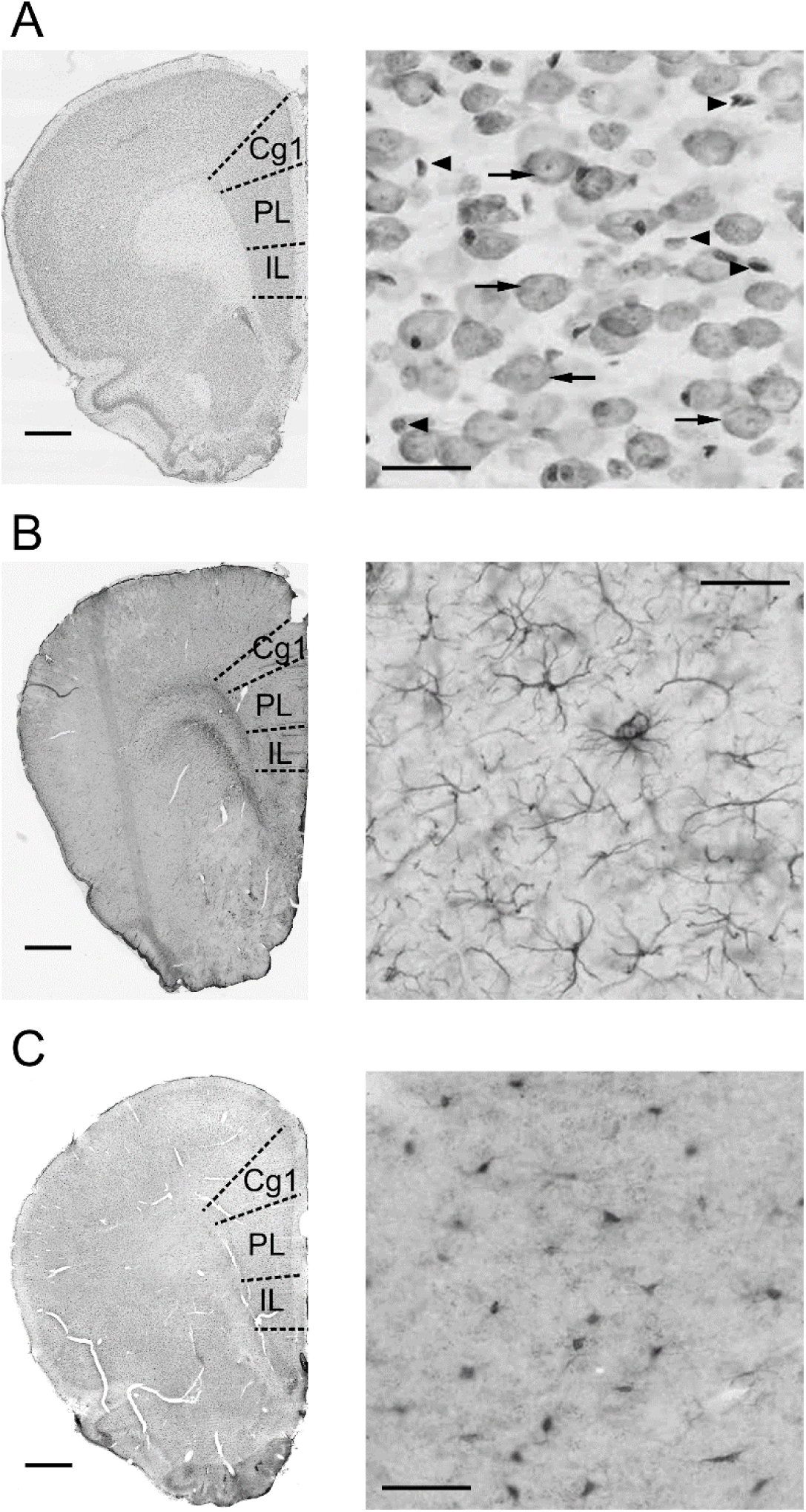
Representative photomicrographs showing examples of CV staining (A) and GFAP (B) and IBA1 immunostaining (C) in the mPFC of control adolescent females. Higher magnification images (right column) of the PL region of the mPFC. The arrows in A indicate CV-stained neuronal cells characterized by pale-stained large euchromatic nuclei with dark-stained nucleoli within the perikaryon. For comparison, the arrowheads in A point to CV-stained nonneuronal cells with strongly stained nuclei and undefined borders between the perikaryon and the nucleus. These types of cells were not included in the analysis. Scale bars: 1 mm (A−C, left column), 30 µm (A, right column) or 50 µm (B, C, right column). Abbreviations: Cg1, cingulate cortex 1; CV, cresyl violet; IL, infralimbic cortex; PL, prelimbic cortex.

Stereological and statistical analyses of regional volume, numerical density and total numbers of neurons and glial cells in the HP revealed that juvenile FLX treatment had no effect on any of the parameters studied in this brain region of adolescent females (ESM_1: Tables S2, S3). Representative photomicrographs showing examples of CV histological staining and GFAP and IBA1 immunostaining in the HP of adolescent females are presented in ESM_2: Fig. S1.

### Effects of juvenile FLX treatment on the mRNA expression of ER stress, apoptosis and synaptic plasticity markers in the mPFC and HP of adolescent females

ER stress, which is strictly related to the unfolded protein response (UPR), is an important cellular process that regulates cell death/survival decisions in response to many different environmental conditions (Hetz & Papa 2018). Statistical analysis revealed that juvenile FLX treatment increased the mRNA levels of all the studied markers of ER stress in the mPFC of adolescent females, including the mRNAs of all the ER stress sensors (Ern1: *t*_10_ = 2.94, *p* = 0.015; Eif2ak3: *t*_10_ = 4.09, *p* = 0.002; Atf6: *t*_10_ = 3.50, *p* = 0.006; Student’s t test) and molecular chaperones belonging to 70-kDa heat shock proteins, such as Hspa5 (*t*_10_ = 3.00, *p* = 0.013) and Hspa1b (*t*_10_ = 2.91, *p* = 0.016; Student’s t test) (Fig. 8A, left panel). Additionally, among the markers of the intrinsic pathway of apoptosis, FLX specifically upregulated the expression of proapoptotic Casp9 and antiapoptotic Bcl2 in the mPFC (Casp 9: *t*_10_ = 2.73, *p* = 0.021; Bcl2: *t*_10_ = 3.66, *p* = 0.004; Casp12: *t*_10_ = 0.273, *p* = 0.790; Casp3: *t*_10_ = 1.04, *p* = 0.322; Bax: *t*_10_ = 0.625, *p* = 0.546, Student’s t test) (Fig. 8A, right panel). However, FLX treatment did not affect the Bax/Bcl2 ratio (VEH: 34.5 ± 5.2, FLX: 29.72 ± 4.33; *t*_10_ = 1.74, *p* = 0.112, Student’s t test).

**Fig. 8.**
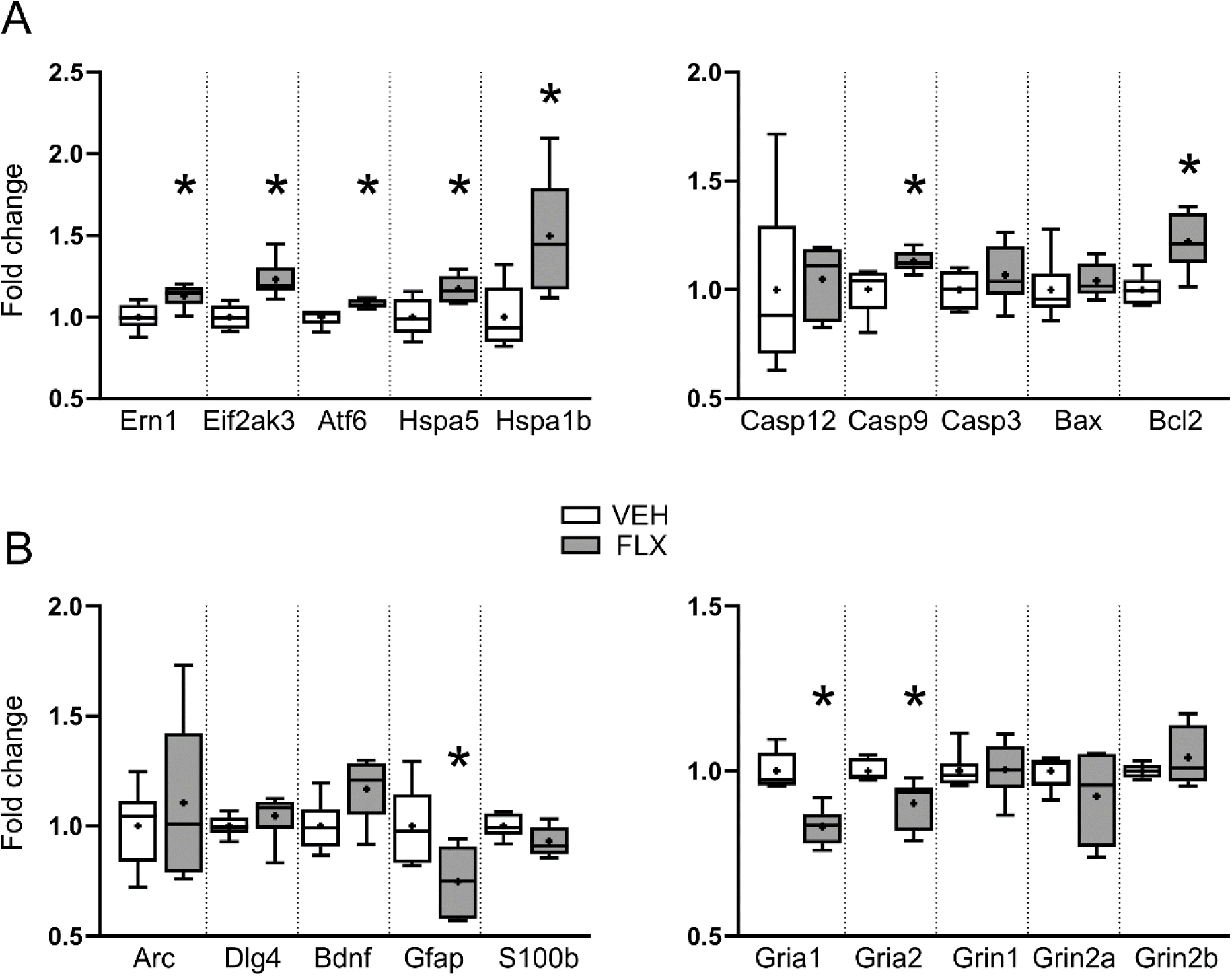
Effects of juvenile FLX treatment on the mRNA expression of ER stress markers (A, left panel), apoptosis markers (A, right panel) and genes involved in synaptic plasticity (B) in the mPFC of adolescent females. The data are presented as box plots and expressed as a fold change relative to the values of VEH rats. In the box plots, the boxes depict the 25th and 75th quartiles, the horizontal line and plus sign correspond to the median and the mean, respectively, whereas the whiskers point to the minimum and maximum of the data (*n* = 6). \**p* < 0.05 compared with VEH rats (Student’s t test in A and B (Gfap, Gria1) and the Mann–Whitney U test in B (Gria2)). Abbreviations: FLX, fluoxetine; VEH, vehicle.

On the other hand, in the mPFC, FLX downregulated the mRNA expression of few genes involved in synaptic plasticity, i.e., the astrocytic marker Gfap (*t*_10_ = 2.47, *p* = 0.033), the AMPA glutamate receptor Gria1 (*t*_10_ = 5.13, *p* = 0.0004, Student’s t test) and Gria2 (*U*_6,6_ = 2.0, *p* = 0.013, Mann–Whitney U test) (Fig. 8B). The mRNA levels of the other synaptic plasticity markers under study were not affected by FLX in the mPFC (Arc: *t*_10_ = 0.612, *p* = 0.554; Dlg4: *U*_6,6_ = 8.0, *p* = 0.128; Bdnf: *t*_10_ = 2.21, *p* = 0.051; S100b: *t*_10_ = 1.94, *p* = 0.084; Grin1: *U*_6,6_ = 14.0, *p* = 0.575; Grin2a: *U*_6,6_ = 16.0, *p* = 0.810; Grin2b: *U*_6,6_ = 16.0, *p* = 0.810) (Fig. 8B).

In the HP of adolescent females, we did not detect any difference in the mRNA expression of ER stress, apoptosis or synaptic plasticity markers between the experimental groups (ESM_1: Table S4, S5).

## Discussion

The goal of our study was to determine whether exposure to FLX during the early-life period, which corresponds to midchildhood and early adolescence in humans, affects the brain maturation processes and behavior of adolescent female rats. We focused on the preadolescence and adolescence periods because during these developmental phases, the brain undergoes substantial structural and functional remodeling and refinement (Selemon 2013). Concurrently, it is also a time of emergence of many psychiatric disorders, such as anxiety and mood disorders, eating disorders and substance use disorders (Paus et al 2008). Consequently, exposure to psychotropic medications, such as FLX, a first-line treatment for many of these disorders in young people, may occur (Paus et al 2008, Selemon 2013). The safety of the administration of FLX during the period of postnatal brain maturation is still questionable.

We found that FLX-treated females presented attenuation of anxiety-like behaviors and impairments in the acquisition and expression of fear memory during adolescence. Juvenile FLX also decreased locomotor activity in response to amphetamine and acute stress (saline injection). Simultaneously, juvenile FLX exposure increased the mPFC volume in all studied subregions and the numbers of neurons, specifically in the Cg1 and PL, as well as the number of astrocytes in the IL. Additionally, FLX-treated females showed higher expression of genes regulating cell survival and reduced mRNA levels of the AMPA glutamate receptors Gria1 and Gria2 in the mPFC. Interestingly, juvenile FLX did not affect any of the studied parameter in the HP of adolescent females.

### Effects of juvenile FLX on the body weight and behavior of adolescent females

In our study, we administered FLX at a dose of 5 mg/kg. This dose was chosen on the basis of many published reports investigating the effects of FLX treatment in juvenile and adolescent rodents (Klomp et al 2012, Klomp et al 2014, Norrholm & Ouimet 2000). Moreover, it corresponds well with the clinically effective dose of FLX, i.e., 20 mg/day (ca. 0.3-0.9 mg/kg) (Beasley et al 2000, Zhou et al 2020), with the assumption that, in rats, drugs are commonly administered at 10-fold higher doses owing to high hepatic drug metabolism and faster elimination (Caccia et al 1990). Our behavioral experiments were started after the 5-day FLX wash-out period to avoid a direct impact of the last FLX injection and to observe the effects over time, i.e., during adolescence. Although we did not measure the plasma levels of FLX and its metabolite norfluoxetine, we assumed that both were not present in the blood of our adolescent females (PND 33-40). FLX elimination *t*_1/2_ is 4–7 h in rats (in humans, 1–4 days), whereas norfluoxetine elimination *t*_1/2_ in rats is approximately 15 h (in humans, approximately 7 days) (Caccia et al 1990). The applied dose and injection schedule of FLX did not affect the body weight of juvenile or adolescent females. Our results are in line with many reports showing that FLX administered during the juvenile and early adolescence periods does not influence weight gain (Homberg et al 2011, Sadegzadeh et al 2020, Wegerer et al 1999). In contrast, exposure to FLX in late adolescence and adulthood results in a decrease in body weight (Aggarwal et al 2016, Homberg et al 2011).

A recently published meta-analysis of animal studies concerning the effects of SSRIs on anxiety-like behavior revealed that this group of drugs generally reduces unconditioned anxiety (Heesbeen et al 2024). However, this analysis did not investigate important characteristics, such as the age of the animals under treatment. Accumulated data clearly show that the effect of FLX on anxiety-like behaviors is age dependent (Kryst et al 2022). Exposure to FLX during juvenile (PNDs 1–21) or broadly defined adolescence (PNDs 20–60) periods usually produces anxiogenic effects when measured in young animals (Adjimann et al 2021, Kryst et al 2022). However, it is worth emphasizing that the majority of studies were performed on males. In contrast, our study revealed that FLX treatment, specifically during PND 20 to PND 28, induced anxiolytic effects in adolescent females. A similar effect was also reported by Sadegzadeh et al. (2020) after the exposure of female rats to FLX at a dose of 5 mg/kg on PND 21-60 (Sadegzadeh et al 2020). Thus, we cannot rule out the possibility of sex-dependent differences in response to FLX in juvenile and adolescent animals.

Numerous studies related to the effects of SSRIs on fear learning have shown that these drugs, including FLX, impair the expression of contextual fear memory in adults but not in adolescents (Heesbeen et al 2023, Norcross et al 2008, Chan et al 2024). However, in the majority of the studies, only male subjects were tested. Interestingly, the present study revealed that juvenile FLX treatment impaired all the studied phases of FC in adolescent females, i.e., the acquisition of FC and expression of both the CFC and AFC. Taken together, our results suggest that juvenile FLX may cause a general inhibition of fear-related emotions in adolescent females (both unconditioned and conditioned). This may underlie the therapeutic action of FLX in patients. However, attenuation of innate and conditioned fear may also reduce the response to negative feedback and lead to an increase in risk-taking behaviors and, consequently, to life-/health-threatening decisions. On the other hand, this can also underlie the phenomenon of emotional blunting, which is frequently observed in patients chronically treated with SSRIs (Marazziti et al 2019).

Potential drug-induced hyper or hypoactivity may affect animal behavior in tests evaluating fear and anxiety and obstruct interpretation of the results. In our study, we did not observe an effect of FLX on the total distance traveled by adolescent females during the light‒dark box test, the acquisition of FC or the novelty-induced locomotion test. However, FLX-treated females displayed greater locomotor activity during the CFC and AFC expression sessions, which may be related to a concurrent reduction in freezing behavior. It should be also taking into account that a modulation of nociception by a drug may affect sensitivity to the foot shock stimulus presented during the session of the acquisition of FC and have nothing to do with fear learning. The 5-HT system is known for its involvement in pain perceptions (Charnay & Leger 2010, Sodhi & Sanders-Bush 2004). Moreover, many studies have shown that acute and chronic FLX treatment has antinociceptive effects that last at least a few hours (Deshmukh 2021, Nayebi et al 2001, Sikka et al 2011). On the other hand, Nelson et al. (1997) reported that repeated FLX injections did not alter foot shock thresholds (Nelson et al 1997). We did not evaluate the effects of FLX on sensory thresholds in our study; however, the 5-day FLX wash-out period before the experiments made potential analgesic activity of the drug less possible under these specific experimental conditions.

Another behavioral characteristic of our adolescent females treated with FLX during juvenility was the blunted response to amphetamine and vehicle injection in the locomotor activity test. Since the 1970s, pharmacological manipulation of the 5-HT system has been known to affect amphetamine-induced hyperlocomotion in adult animals. Specifically, 5-HT blocking agents were found to potentiate the locomotor response to this psychostimulant, whereas 5-HT agonists, especially when injected into the nucleus accumbens, reduced such behavior (Carter & Pycock 1978, Hollister et al 1976). However, extensive data collected over the years have revealed that acute and chronic FLX treatment, known to increase 5-HT levels, potentiates hyperlocomotion induced by amphetamine for up to 72 h (3 days) but not 5 days after the withdrawal of FLX treatment, when no change in response to amphetamine has been observed (Dziedzicka-Wasylewska et al 2002, Sills et al 1999, Sills et al 2000). Because FLX simultaneously and transiently increases brain amphetamine levels, this phenomenon was interpreted as the inhibitory effect of FLX on amphetamine metabolism, i.e., the pharmacokinetic effect, and not as a direct or adaptive interaction between the 5-HT system and the mesolimbic dopaminergic system (Sills et al 1999, Sills et al 2000). Nevertheless, in our study, which was conducted in juveniles, during the critical period of brain maturation, we observed not only a lack of changes in amphetamine-induced hyperlocomotion but also a decrease in hyperlocomotion 5 days after the withdrawal of FLX. A similar effect, i.e., a blunted response to amphetamine together with reduced motivation and dopaminergic activation in a sucrose reward, was reported by Menezes et al. (2021) in adult mice treated with FLX during the perinatal period (PNDs 2-11) (Menezes et al 2021). Taken together, these findings and our data suggest that high levels of 5-HT during the early postnatal period may affect the developmental trajectory and function of the dopaminergic system. The dopaminergic system develops later than the 5-HT system and is therefore very sensitive to early-life perturbations (Kalsbeek et al 1988).

Interestingly, FLX-treated adolescent females also presented lower locomotor activity in response to vehicle injection (acute stress) but not to novelty in general. These findings may suggest that juvenile FLX exposure induced disturbances in both the mesolimbic and mesocortical dopaminergic systems. Both FLX and acute stress cause the release of dopamine in the mPFC (Butts et al 2011, Bymaster et al 2002). Whereas, the mesocortical dopaminergic pathway exerts inhibitory control on dopamine release in the nucleus accumbens and, consequently, may affect locomotor activity (Pascucci et al 2007, Tzschentke 2001). In summary, these data imply that juvenile FLX treatment may have an inhibitory effect on the dopaminergic reward system (Flores-Ramirez et al 2019, Lee & Lee 2012), which may have positive consequences in the context of high vulnerability to different kinds of addictions observed during adolescence (Paus et al 2008). On the other hand, such a phenomenon may also potentially underlie the emotional blunting and reduced responsiveness to positive and negative feedback and stimuli observed in some patients treated with SSRIs (Marazziti et al 2019).

### Effects of juvenile FLX on the maturation of the mPFC and HP in adolescent females

The next goal of our study was to assess the effects of juvenile FLX exposure on several aspects of brain maturation, with a specific focus on two brain regions, e.g., the mPFC and HP. The mPFC and HP are engaged in higher cognitive function and in the expression of both conditioned and unconditioned fear (Bertagna et al 2021, Ghasemi et al 2022, Lisboa et al 2010, Sierra-Mercado et al 2011). These brain regions are also involved in the pathophysiology of mood and anxiety disorders (Drevets et al 2008, Myers-Schulz & Koenigs 2012), which are highly prevalent and often cooccur in childhood and adolescence (Paus et al 2008). Our study revealed strong region-specific effects of FLX on the studied parameters, which were present exclusively in the mPFC. This observation is in accordance with the fact that the mPFC has a uniquely prolonged developmental trajectory and is one of the last brain regions to mature (Caballero et al 2016, Mills et al 2014). This makes the mPFC especially vulnerable to environmental insults. We observed that FLX-treated adolescent females generally presented greater mPFC volume and greater numbers of neurons than control females did, specifically in Cg1 and PL regions of the mPFC. Simultaneously, the numerical density of the neurons did not differ between the experimental groups. These results indicate that juvenile FLX delays neurodevelopmental apoptosis and presumably synapse pruning, two crucial processes that occur during adolescence in the mPFC (Brenhouse & Andersen 2011, Selemon 2013). 5-HT is known as a regulator of programmed cell death during development (Sodhi & Sanders-Bush 2004). High levels of 5-HT due to 5-HTT knockout have been shown to reduce apoptosis (Altamura et al 2007, Persico et al 2003), as in our study, with transient inhibition of 5-HTT by FLX. Additionally, specific polymorphisms in the 5-HTT promoter region (5-HTTLPR S vs. L), which determine its transcriptional activity and regulate 5-HTT expression, affect gray matter density in humans (Canli et al 2005).

Analysis of the mRNA levels of apoptosis markers in the mPFC revealed increased expression of the antiapoptotic protein Bcl2 in FLX-treated adolescent females. On the other hand, we also detected increased levels of the proapoptotic Casp9, which can potentially be explained as compensatory or counteractive effects. Interestingly, we also observed FLX-induced upregulation of genes involved in ER stress and UPR processes. These processes are known to influence cell death/survival decisions in response to a wide range of environmental conditions, which ultimately results in disruption of protein homeostasis and accumulation of misfolded proteins (Almanza et al 2019, Hetz & Papa 2018). ER stress is sensed by three ER transmembrane proteins, i.e., inositol-requiring enzyme 1 (IRE1α), RNA-activated protein kinase (PKR)-like endoplasmic reticulum kinase (PERK) and activating transcription factor 6 (ATF6) encoded by the Ern1, Eif2ak3 and Atf6 genes, respectively. Under physiological conditions, these three sensors are bound to HSPA5. When ER stress is induced, HSPA5 dissociates from sensor proteins and facilitates their activation and progression of the UPR. In general, the UPR is a protective process that leads to attenuation of general translation, chaperones upregulation and elimination of misfolded proteins. However, if the prosurvival potential of a cell is depleted, programmed cell death pathways are induced through the proapoptotic component of the UPR (Almanza et al 2019, Hetz & Papa 2018). The UPR and ER stress have been implicated in the pathophysiology of many different diseases, including mental disorders, such as mood and anxiety disorders (Almanza et al 2019, Hetz & Papa 2018). Increased expression of ER stress and UPR markers has been observed in animal models of depression and anxiety (Le-Niculescu et al 2011, Solarz et al 2021b, Tang et al 2018, Yang et al 2018), as well as in patients with major depressive disorder and posttraumatic stress disorder (in leucocytes and the mPFC) (Nevell et al 2014, Yoshino & Dwivedi 2020). Therefore, it is not surprising that ER stress and the UPR are considered potential targets of antidepressants, especially SSRIs (Jozwiak-Bebenista et al 2022, Yang et al 2018). Our study confirms this hypothesis.

Interestingly, we also found that juvenile FLX affected glial cells. Specifically, we observed that FLX-treated females presented an increased number of astrocytes and a trend toward a reduced number of microglia in the IL. These results imply that juvenile exposure to FLX interferes with the developmental processes of glial proliferation and/or apoptosis, which are usually intensified during the first three weeks of postnatal life in rodents (Bandeira et al 2009, Nikodemova et al 2015). The simultaneous decrease in the mRNA levels of Gfap observed in our study rather excludes the possibility that the FLX-induced increase in the number of GFAP-IR astrocytes results from enhanced astrocyte reactivity. Our results are in line with a large amount of accumulating data showing that, in addition to neurons, antidepressants may directly act on glial cells (Czeh & Di Benedetto 2013, Kinoshita et al 2018). Glial pathology is observed in mood and anxiety disorders (Verkhratsky et al 2023). Glial cells express 5-HTT and 5-HT receptors (Inazu et al 2001) (Albertini et al 2023, Verkhratsky et al 2023). FLX has been shown to increase nonneuronal cell proliferation (Czeh et al 2007, Kodama et al 2004) and gliotransmission of ATP, which may underlie its therapeutic action (Kinoshita et al 2018). However, during sensitive periods of brain development, the consequences of such FLX action are difficult to predict (Albertini et al 2023).

During early development, 5-HT plays a crucial role not only in neuronal circuit formation but also in synaptic plasticity (Kraus et al 2017, Sodhi & Sanders-Bush 2004). Modulation of synaptic plasticity is considered an important mechanism of the therapeutic action of antidepressants, including SSRIs (Duman 2004). Neuroplasticity underlies learning, memory and adaptation processes. AMPA glutamate receptors are among the key players involved in synaptic plasticity. Changes in the availability of postsynaptic AMPA receptors shape most forms of functional synaptic plasticity, i.e., long-term potentiation (LTP), long-term depression (LTD) and homeostatic scaling (Diering & Huganir 2018). Reduced expression of AMPA receptors has been observed postmortem in the brains of patients suffering from mood disorders (Beneyto et al 2007, Duric et al 2013). Antidepressants, including SSRIs, are known to upregulate AMPA receptor expression in adult animals (Barbon et al 2011, Gerace et al 2023). In contrast, our study revealed that juvenile FLX decreased the mRNA levels of Gria1 and Gria2 (encoding the AMPA receptors GluA1 and GluA2, respectively) in adolescent females, suggesting that FLX has a developmental time- and/or sex-specific effects on synaptic plasticity.

### Limitations of the study

The major limitation of our present study was that male subjects were not included in the experiments. However, epidemiological data clearly show that females have a significantly greater prevalence of anxiety, depression and eating disorders than males do (Breslau et al 2017, Hornberger et al 2021, Lewis et al 2020). It is believed that sex differences in the prevalence of the abovementioned diseases originate during childhood and increase during adolescence (after puberty in the case of depression) (Breslau et al 2017, Jacobs 2009, Lewis et al 2020). All of these disorders are characterized by an imbalance in 5-HT neurotransmission, and SSRIs are considered the first-line pharmacological treatment for both depression and anxiety in young people (Jacobs 2009, Rapee et al 2023, Zhou et al 2020). Regardless, there are few preclinical studies performed on females aimed at studying the effects of SSRIs on brain development and function. It is worth emphasizing that, preadolescence/adolescence is a time of intensified divergence between males and females in the trajectory of brain development, physiology and behavior (Giedd et al 2012, Lenroot & Giedd 2010). Therefore, a broad range, comprehensive studies, simultaneously on males and females, with different schedules of FLX treatment and time points of data analyses are needed to properly evaluate sex differences in response to juvenile FLX exposure. Nevertheless, this was beyond the scope of our preliminary study.

## Conclusions

In summary, juvenile FLX treatment influenced important processes during early postnatal maturation of the mPFC, such as developmental apoptosis and/or glial proliferation. On the basis of the changes observed in mPFC volume and gene expression, FLX likely also influenced synapse pruning and synaptic plasticity. These results suggest that juvenile exposure to FLX may affect proper neuronal circuit formation and function and, consequently, produce specific behavioral phenotypes, such as the inhibition of fear-related emotions and a blunted response to reward or acute stress in adolescent females. The ultimate repercussions of early-life FLX are difficult to predict; therefore, this psychotropic drug should be used with particular caution in young people.

## Supporting information

ESM_1 Tables S1_S5

ESM_2 FigS1

## List of Abbreviations

AMPA: α-amino-3-hydroxy-5-methyl-4-isoxazolepropionic acid
AFC: Auditory fear conditioning
ATF6: Activating transcription factor 6
CFC: Contextual fear conditioning
Cg1: Cingulate cortex 1
CV: Cresyl violet
DG: Dentate gyrus
Eif2ak3: Eukaryotic translation initiation factor 2 α kinase 3
alias: PERK: RNA-activated protein kinase **(PRK)**-like endoplasmic reticulum kinase
ER: Endoplasmic reticulum
Ern1: Endoplasmic reticulum to nucleus signaling 1
alias: IRE1α: Inositol-requiring enzyme 1
FC: Fear conditioning
FLX: Fluoxetine
GFAP: Glial fibrillary acidic protein
HP: Hippocampus
Hspa5: Heat shock protein family A member 5
alias: GRP78: glucose-regulated protein 78 kDa
Hspa1b: Heat shock protein family A member 1b
5-HT: 5-hydroxytryptamine
5-HTT: Serotonin transporter
IBA1: Ionized calcium-binding adapter molecule
IL: Infralimbic cortex
IR: Immunoreactive
mPFC: Medial prefrontal cortex
PL: Prelimbic cortex
PND: Postnatal day
SSRIs: Selective serotonin reuptake inhibitors
UPR: Unfolded protein response
VEH: Vehicle

## CrediT authorship contribution statement

Joanna Kryst: Investigation, Methodology, Data curation, Visualization; Agnieszka Chocyk: Conceptualization, Methodology, Formal analysis, Visualization, Writing – Original Draft, Project administration, Funding acquisition; Anna Solarz-Andrzejewska: Investigation, Methodology, Visualization; Iwona Majcher-Maślanka: Investigation, Methodology.

## Conflict of interest

The authors declare that the research was conducted in the absence of any commercial or financial relationships that could be construed as a potential conflict of interest.

## Acknowledgements

This work was supported by grant Opus 2017/25/B/NZ7/00174 from the National Science Centre, Poland to A.C. and by the statutory activity of the Maj Institute of Pharmacology, Polish Academy of Sciences, Smętna Street 12, 31-343 Kraków, Poland.

## Notes

### Competing Interest Statement

The authors have declared no competing interest.

