## Supplementary material for "Juvenile fluoxetine treatment affects the maturation of the medial prefrontal cortex and behavior of adolescent female rats": ESM_1 Tables S1_S5

Table S1. Effects of juvenile FLX treatment on the numerical density of CV-positive neurons, GFAP-IR astrocytes and IBA1-IR microglial cells in the mPFC of adolescent female rats

| Cell marker | mPFC region | Group | Cell density/mm$^3$ | Statistic |
|---|---|---|---|---|
| CV | Cg1 | VEH | 132 151.59 ± 9992.97 | $t_8 = 0.531$, $p = 0.610$ |
| | | FLX | 135 162.18 ± 7798.08 | |
| | PL | VEH | 175 831.44 ± 9246.12 | $t_8 = 0.132$, $p = 0.898$ |
| | | FLX | 175 140.08 ± 7159.45 | |
| | IL | VEH | 190 147.14 ± 17 182.14 | $t_8 = 1.001$, $p = 0.346$ |
| | | FLX | 180 694.06 ± 12 291.14 | |
| GFAP | Cg1 | VEH | 61 174.01 ± 8949,01 | $t_8 = 0.876$, $p = 0.406$ |
| | | FLX | 57 224.89 ± 4635.47 | |
| | PL | VEH | 71 657.21 ± 11 547.46 | $t_8 = 1.351$, $p = 0.214$ |
| | | FLX | 63 612.08 ± 6638.77 | |
| | IL | VEH | 73 361.32 ± 8337.70 | $t_8 = 0.911$ $p = 0.389$ |
| | | FLX | 68 229.28 ± 9435.24 | |
| IBA1 | Cg1 | VEH | 26 390.76 ± 897.93 | $t_8 = 1.029$, $p = 0.333$ |
| | | FLX | 25 375.13 ± 2014.99 | |
| | PL | VEH | 27 364.52 ± 863.57 | $t_8 = 1.807$, $p = 0.108$ |
| | | FLX | 25 924.55 ± 1558.34 | |
| | IL | VEH | 26 001.56 ± 1600.15 | $t_8 = 2.215$, $p = 0.058$ |
| | | FLX | 24.123.74 ± 1016.59 | |

Data indicate the numerical density of cells in the subregions of the mPFC estimated by stereological method (the mean ± SD, $n = 5$). Student's t-test did not show any statistically significant differences between treatment groups. Abbreviations: Cg1, cingulate cortex 1; FLX, fluoxetine; GFAP; glial fibrillary acidic protein; IBA1, ionized calcium-binding adaptor molecule 1; IL, infralimbic cortex; IR, immunoreactive; PL, prelimbic cortex; VEH, vehicle.

Table S2. Effects of juvenile FLX treatment on HP volume in adolescent female rats

| **Region** | **Group** | **Volume (mm$^3$)** | **Statistic** |
|---|---|---|---|
| CA1 | VEH<br>FLX | 22.26 ± 2.20<br>23.13 ± 2.97 | $t_8 = 0.524$, $p = 0.615$ |
| CA2-CA3 | VEH<br>FLX | 15.56 (1.19)<br>15.77 (3.38) | $U_{5,5} = 9.00$, $p = 0.531$ |
| DG | VEH<br>FLX | 16.20 (1.50)<br>17.37 (0.11) | $U_{5,5} = 9.00$, $p = 0.531$ |

The data are presented as the mean ± SD or median (IQR), $n = 5$. Statistical analysis (Student's t-test or Mann–Whitney U test, respectively) did not show any significant differences between treatment groups. Abbreviations: CA1-CA3, cornu Ammonis fields; DG, dentate gyrus; FLX, fluoxetine; IQR, interquartile range; VEH, vehicle.

Table S3. Effects of juvenile FLX treatment on density and the estimated total number of CV-positive neurons, GFAP-IR astrocytes and IBA1-IR microglial cells in the HP of adolescent female rats

| **Cell marker** | **HP region** | **Parameter** | **Group** | **Numerical values** | **Statistic** |
|---|---|---|---|---|---|
| CV | CA1 | Cell density/mm$^3$ | VEH<br>FLX | 55 417.96 ± 3684.04<br>56 630.45 ± 5321.61 | $t_8 = 0.419$, $p = 0.686$ |
| | | Cell no./region | VEH<br>FLX | 1 257 739.80 ± 83 611.31<br>1 285 258.03 ± 120 776.69 | |
| | CA2-CA3 | Cell density/mm$^3$ | VEH<br>FLX | 67 963.25 (10 749.76)<br>66 387.11 (3817.16) | $U_{5,5} = 6.00$, $p = 0.210$ |
| | | Cell no./region | VEH<br>FLX | 1 102 187.09 (174 333.15)<br>1 076 626.30 (61 904.37) | |
| | DG | Cell density/mm$^3$ | VEH<br>FLX | 95 815.39 ± 9660.77<br>89 788,05 ± 9029.63 | $t_8 = 1.019$, $p = 0.338$ |
| | | Cell no./region | VEH<br>FLX | 1 547 127.93 ± 155 992.16<br>1 449 804.57 ± 145 801.06 | |
| GFAP | CA1 | Cell density/mm$^3$ | VEH<br>FLX | 57 618.87 (6776.91)<br>55 860.51 (15 661.88) | $U_{5,5} = 11.00$, $p = 0.834$ |
| | | Cell no./region | VEH<br>FLX | 1 069 997.48 (125 849.02)<br>1 037 344.19 (290 845.27) | |
| | CA2-CA3 | Cell density/mm$^3$ | VEH<br>FLX | 59 072.37 ± 6634.69<br>58 587.79 ± 7071.06 | $t_8 = 0.112$, $p = 0.914$ |
| | | Cell no./region | VEH<br>FLX | 911 075.87 ± 102 327.05<br>903 602.19 ± 109 057.22 | |
| | DG | Cell density/mm$^3$ | VEH<br>FLX | 54 396.92 ± 7121.13<br>53 680.79 ± 10 077.43 | $t_8 = 0.130$, $p = 0.900$ |
| | | Cell no./region | VEH<br>FLX | 765 651.61 ± 100 231.87<br>755 571.90 ± 141 842.55 | |
| IBA1 | CA1 | Cell density/mm$^3$ | VEH<br>FLX | 31 690.23 ± 2348.37<br>29 675.48 ± 1510.57 | $t_8 = 1.613$, $p = 0.145$ |
| | | Cell no./region | VEH<br>FLX | 564 144.17 ± 41 805.23<br>528 277.82 ± 26 890.92 | |
| | CA2-CA3 | Cell density/mm$^3$ | VEH<br>FLX | 27 250.64 ± 2148.55<br>25 907.65 ± 1162.25 | $t_8 = 1.229$, $p = 0.254$ |
| | | Cell no./region | VEH<br>FLX | 407 872.25 ± 32 158.29<br>387 771.14 ± 17 395.91 | |
| | DG | Cell density/mm$^3$ | VEH<br>FLX | 30 564.53 (220.36)<br>27 029.48 (610.70) | $U_{5,5} = 5.00$, $p = 0.144$ |
| | | Cell no./region | VEH<br>FLX | 395 690.85 (2852.80)<br>349 925.78 (7906.12) | |

Data indicate the numerical density and the total numbers of cells per region estimated by stereological method (the mean ± SD or median (IQR), $n$ = 5). Statistical analysis (Student's t-test or Mann–Whitney U test, respectively) did not show any significant differences between treatment groups. Abbreviations: CA1-CA3, cornu Ammonis fields; CV, cresyl violet; DG, dentate gyrus; FLX, fluoxetine; GFAP, glial fibrillary acidic protein; IBA1, ionized calcium-binding adaptor molecule 1; IQR, interquartile range; IR, immunoreactive; VEH, vehicle.

Table S4. Effects of juvenile FLX treatment on mRNA expression of ER stress, UPR and apoptotic markers in the HP of adolescent female rats

| **Gene** | **Group** | **Relative mRNA level** | **Statistic** |
|---|---|---|---|
| Hspa5 | VEH<br>FLX | 0.205360 ± 0.025054<br>0.209877 ± 0.025486 | $t_{10}$ = 0.310, $p$ = 0.763 |
| Hspa1b | VEH<br>FLX | 0.000055 (0.000015)<br>0.000058 (0.000006) | $U_{6,6}$ = 15.00, $p$ = 0.689 |
| Ern1 | VEH<br>FLX | 0.004668 ± 0.000388<br>0.004554 ± 0.000369 | $t_{10}$ = 0.521, $p$ = 0.614 |
| Eif2ak3 | VEH<br>FLX | 0.025133 (0.008932)<br>0.023569 (0.000422) | $U_{6,6}$ = 13.00, $p$ = 0.471 |
| Atf6 | VEH<br>FLX | 0.031751 ± 0.000789<br>0.030892 ± 0.001048 | $t_{10}$ = 1.605, $p$ = 0.139 |
| Casp12 | VEH<br>FLX | 0.000234 ± 0.000061<br>0.000211 ± 0.000023 | $t_{10}$ = 0.894, $p$ = 0.392 |
| Casp9 | VEH<br>FLX | 0.014307 ± 0.002274<br>0.015655 ± 0.001480 | $t_{10}$ = 1.217, $p$ = 0.252 |
| Casp3 | VEH<br>FLX | 0.002428 ± 0.000207<br>0.002301 ± 0.000099 | $t_{10}$ = 1.363, $p$ = 0.203 |
| Bax | VEH<br>FLX | 0.054923 ± 0.002765<br>0.050959 ± 0.003192 | $t_{10}$ = 2.300, $p$ = 0.054 |
| Bcl2 | VEH<br>FLX | 0.002598 (0.000214)<br>0.002649 (0.000487) | $U_{6,6}$ = 17.00, $p$ = 0.936 |

The mRNA expression was determined by RT-qPCR and presented as relative values of mRNA levels in arbitrary units. The data are presented as the mean ± SD or median (IQR), $n$ = 6. Statistical analysis (Student's t-test or Mann–Whitney U test, respectively) did not show any significant differences between treatment groups. Abbreviations: Atf6, activating transcription factor 6; Casp, caspase; Eif2ak3, eukaryotic translation initiation factor 2 α kinase 3; ER, endoplasmic reticulum; Ern1, endoplasmic reticulum to nucleus signaling 1; FLX, fluoxetine; Hspa, heat shock protein family A members (1b and 5); IQR, interquartile range; UPR, unfolded protein response; VEH, vehicle.

Table S5. Effects of juvenile FLX treatment on mRNA expression of synaptic plasticity markers in the HP of adolescent female rats

| **Gene** | **Group** | **Relative mRNA level** | **Statistic** |
|---|---|---|---|
| Arc | VEH | 0.051108 ± 0.008370 | $t_{10}$ = 0.515, $p$ = 0.618 |
| | FLX | 0.048726 ± 0.007637 | |
| Dlg4 | VEH | 0.309229 ± 0.030950 | $t_{10}$ = 0.073, $p$ = 0.943 |
| | FLX | 0.310401 ± 0.024287 | |
| Bdnf | VEH | 0.008803 (0.000756) | $U_{6,6}$ = 17.00, $p$ = 0.936 |
| | FLX | 0.008306 (0.001205) | |
| Gfap | VEH | 0.236442 (0.023216) | $U_{6,6}$ = 13.00, $p$ = 0.471 |
| | FLX | 0.261544 (0.020841) | |
| S100b | VEH | 0.231581 ± 0.010891 | $t_{10}$ = 1.801, $p$ = 0.102 |
| | FLX | 0.220180 ± 0.011042 | |
| Gria1 | VEH | 0.216458 ± 0.018758 | $t_{10}$ = 1.340, $p$ = 0.210 |
| | FLX | 0.228923 ± 0.012947 | |
| Gria2 | VEH | 0.213979 ± 0.016093 | $t_{10}$ = 1.307, $p$ = 0.221 |
| | FLX | 0.225104 ± 0.013257 | |
| Grin1 | VEH | 0.256103 ± 0.014286 | $t_{10}$ = 0.496, $p$ = 0.631 |
| | FLX | 0.259662 ± 0.010230 | |
| Grin2a | VEH | 0.194057 (0.007946) | $U_{6,6}$ = 6.00, $p$ = 0.066 |
| | FLX | 0.198850 (0.004116) | |
| Grin2b | VEH | 0.121905 ± 0.008363 | $t_{10}$ = 0.674, $p$ = 0.515 |
| | FLX | 0.125420 ± 0.009654 | |

The mRNA expression was determined by RT-qPCR and presented as relative values of mRNA levels in arbitrary units. The data are presented as the mean ± SD or median (IQR), $n$ = 6. Statistical analysis (Student’s t-test or Mann–Whitney U test, respectively) did not show any significant differences between treatment groups. Abbreviations: Arc, activity-regulated cytoskeleton-associated protein; Bdnf, brain-derived neurotrophic factor; Dlg4, discs large homolog 4; FLX, fluoxetine; Gfap, glial fibrillary acidic protein; Gria1 and 2, glutamate ionotropic receptor AMPA type, subunit 1 and 2; Grin1, 2a and 2b, glutamate ionotropic receptor NMDA type, subunit 1, 2a and 2b; IQR, interquartile range; S100b, calcium-binding protein B; VEH, vehicle.
