## Supplementary material for "Juvenile fluoxetine treatment affects the maturation of the medial prefrontal cortex and behavior of adolescent female rats": ESM_2 FigS1

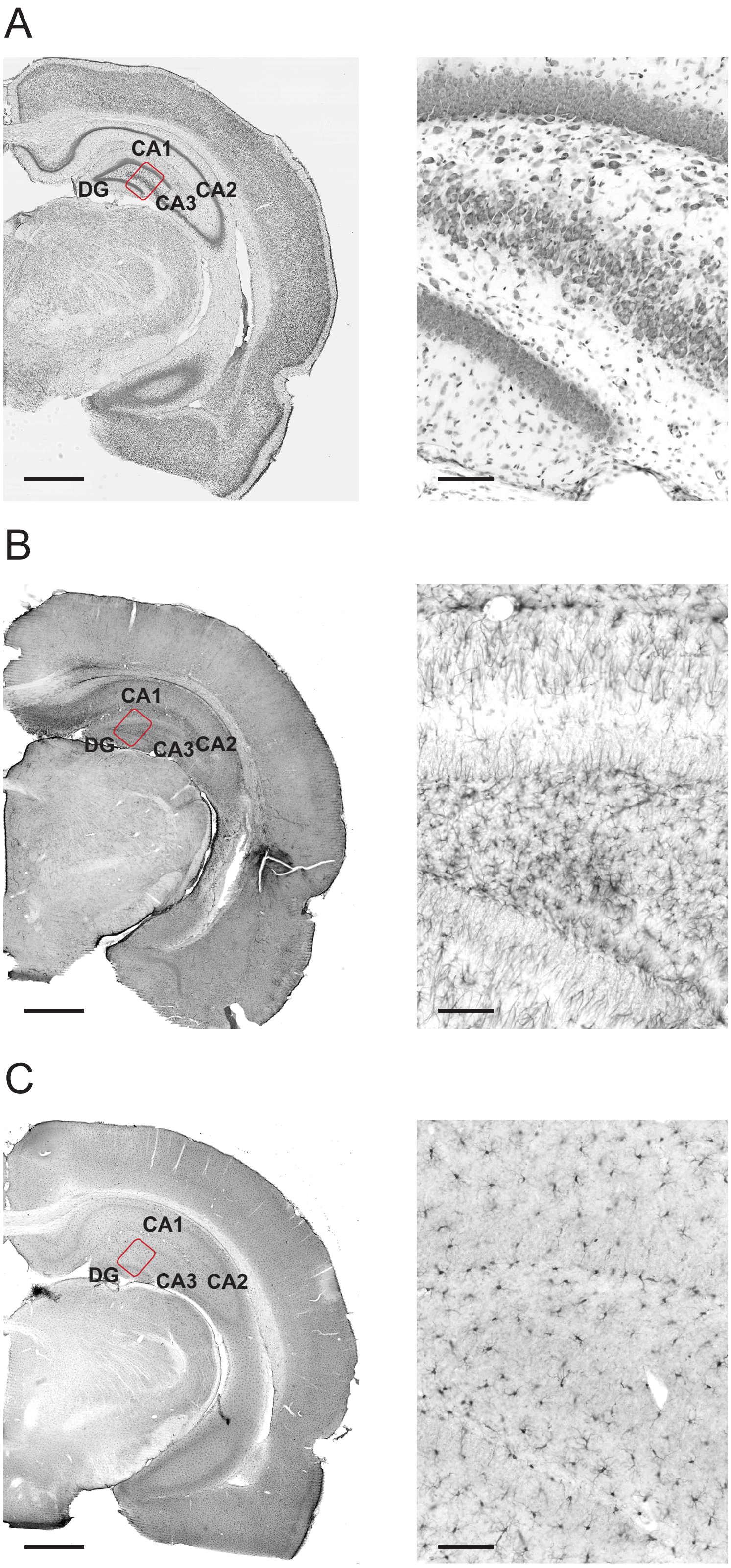


Fig. S1. Representative photomicrographs showing examples of CV staining (A), GFAP (B), and IBA1 immunostaining (C) in the HP of control adolescent females. Higher magnification images (right column) come from the region indicated by red rectangles (left column). Scale bars: 1 mm (A-C, left column), 100 µm. (A-C, right column). Abbreviations: CA1-CA3, cornu Ammonis fields; CV, cresyl violet; DG, dentate gyrus.
